# Variant-to-gene mapping in ocular cell types identifies candidate effector genes for primary open-angle glaucoma

**DOI:** 10.64898/2026.02.20.707051

**Authors:** Vrathasha Vrathasha, Matthew C. Pahl, James A. Pippin, Sergei Nikonov, Jie He, Mina Halimitabrizi, Laxmi Moksha, Rebecca Salowe, Amy-Ann Edziah, Yuki Bradford, Yan Zhu, Harini V. Gudiseva, Venkata R. M. Chavali, Bruna Lopes da Costa, Anne Marie Berry, Peter M. J. Quinn, Qi N. Cui, Eydie Miller-Ellis, Prithvi S. Sankar, Ahmara G. Ross, Victoria Addis, Shefali S. Verma, Andrew D. Wells, Struan F. A. Grant, Joan M. O’Brien

**Affiliations:** Center for Genetics of Complex Disease, Department of Ophthalmology, University of Pennsylvania, Philadelphia, PA 19104, USA; F.M. Kirby Center for Molecular Ophthalmology, University of Pennsylvania, Philadelphia, PA 19104, USA; Center for Spatial and Functional Genomics, The Children’s Hospital of Philadelphia, Philadelphia, PA 19104, USA; Division of Genetic and Genomic Medicine, The Children’s Hospital of Philadelphia, Philadelphia, PA 19104, USA; Department of Medicine, Perelman School of Medicine, University of Pennsylvania, Philadelphia, PA 19104, USA; Department of Pathology and Laboratory Medicine, Perelman School of Medicine, University of Pennsylvania, Philadelphia, PA, USA; Department of Pathology, The Children’s Hospital of Philadelphia, Philadelphia, PA 19104, USA; Institute for Immunology & Immune Health, Perelman School of Medicine, University of Pennsylvania, Philadelphia, PA 19104, USA; Department of Pediatrics, Perelman School of Medicine, University of Pennsylvania, Philadelphia, PA 19104, USA; Department of Genetics, Perelman School of Medicine, University of Pennsylvania, Philadelphia, PA 19104, USA; Division of Diabetes and Endocrinology, The Children’s Hospital of Philadelphia, 3615 Civic Center Boulevard, Philadelphia, PA 19104, USA

## Abstract

Primary open-angle glaucoma (POAG), a leading cause of irreversible blindness worldwide, has a strong genetic basis. The Primary Open-Angle African Ancestry Glaucoma Genetics study previously identified 46 risk loci associated with this disease. To pinpoint causal variants and their corresponding effector genes, we analyzed gene expression, chromatin accessibility, and conformation in two ocular cell types: trabecular meshwork cells (hTMCs) and retinal ganglion cells derived from induced pluripotent stem cells (hiPSC-RGCs). We identified 24 candidate genes in hTMCs and 56 in hiPSC-RGCs. Several prioritized genes showed altered mRNA expression, implicating pathways involved in oxidative stress, tissue homeostasis, neuronal maintenance, and retinal degeneration. Among these genes, *ARHGEF12* was selected for further validation because local and distal promoter interactions nominated it in both cell types, and prior evidence reproducibly links it to POAG. While its role in hTMCs is established, its function in RGCs is unclear. hiPSC-RGCs generated from a POAG donor homozygous for the risk allele exhibited reduced *ARHGEF12* expression, altered morphology, and disrupted neuronal activity. By linking POAG risk loci to candidate effector genes in ocular cell types, this study provides a framework for functional evaluation of additional POAG risk variants.

## Introduction

Over the past two decades, genome-wide association studies (GWAS) have revealed thousands of single-nucleotide polymorphisms (SNPs) that are statistically associated with complex phenotypes and disease risk.^1–3^ The vast majority of these GWAS-implicated signals are located in non-coding regions of the genome, presumably influencing disease susceptibility by altering the expression of one or more target genes.^4–6^ However, the assumption that the nearest gene to the GWAS-implicated signal represents the true effector gene has proven insufficient. Instead, effector genes can be located tens or hundreds of kilobases distal to regulatory elements due to long-range interactions at the 3D genome level.^4,6–12^ Because GWAS does not unambiguously reveal the target genes or cell types affected by the discovered signals, this complexity hinders the interpretation of potential disease mechanisms.^8,13–15^

Historically, the study of 3D genome organization has been challenging due to a lack of relevant methodologies. However, over the past decade, chromosome conformation capture (3C)-based techniques, such as Hi-C and its derivatives, have enabled mapping of putative contacts across a range of genomic distances.^16,17^ Structural genomic approaches have been successfully applied to several complex genetic disorders, implicating novel risk genes. For example, for type 2 diabetes, 3D genomic approaches revealed that a causal variant for the strongest signal in the *TCF7L2* gene^18,19^ also functions as a regulatory region for *ACSL5*.^20^ Similar approaches in osteoporosis^6^, obesity^21–23^, carpal tunnel syndrome^8^, type 2 diabetes^24^, and rheumatoid arthritis^25^ have also implicated novel causal target genes that drove functional follow-up studies.

However, such structural genomic efforts in the context of primary open-angle glaucoma (POAG) remain limited,^10,26^ despite the disease being a leading cause of irreversible blindness worldwide.^27–29^ POAG has a strong genetic component, with heritability estimates ranging from 13% to 81%.^30–33^ Several monogenic mutations have been reported for POAG, but only account for 5% of all cases.^34–37^ Thus far, GWAS analyses have identified 329 risk loci associated with POAG and its related endophenotypes.^38–54^ However, the vast majority of these loci are harbored in non-coding areas of the genome and thus do not directly pinpoint the underlying specific gene(s), cell types, or mechanisms.^40^ Translating these results into biological insights requires systematic identification of putative causal variants and the corresponding candidate effector genes, together with functional verification in either cellular or animal models.^24,55–57^ Indeed, Wang *et al* reported that using scATAC-seq-derived peaks in the human retina could assign a SNP (rs4821699), located in the intronic region of *TRIOBP,* as a contributing causal variant to glaucoma by altering the gene’s expression in retinal ganglion cells (RGCs).^12^ Additionally, the study conducted by Hamel *et al* has prioritized causal regulatory mechanisms, target genes, and cell types using expression and splicing QTLs, retinal Hi-C, and single-cell expression data from the outflow pathways, retina, and optic nerve head.^58^ Nevertheless, a comprehensive variant-to-gene map of POAG risk loci in disease-relevant ocular cell types remains lacking, limiting mechanistic interpretation of genetic associations and prioritization of candidate effector genes for functional study.

The Primary Open-Angle African Ancestry Glaucoma Genetics (POAAGG) study previously conducted a GWAS on POAG in African ancestry individuals, identifying 46 risk loci.^38^ Although individuals of African ancestry are four to five times more likely to be affected by the disease and up to 15 times more likely to go blind, only about 2% of all genetic studies and GWAS’s are conducted in this particular population.^59,60^ Although genetic regulation is often shared across populations,^61^ ancestry-specific differences have been observed, as shown by expression QTLs in admixed and African samples compared to European tissue samples in GTEx.^62^ Thus, in this study, we employed structural genomic approaches to implicate causal variants and their corresponding effector genes at the POAG loci in cell lines derived from African ancestry individuals. We generated 3D genomics data using two glaucoma-relevant ocular cell lines: human trabecular meshwork cells (hTMCs) and retinal ganglion cells derived from human induced pluripotent stem cells (hiPSC-RGCs). Our analyses implicated 24 genes in hTMCs and 56 genes in hiPSC-RGCs. Expression analyses of prioritized candidates revealed altered regulation of genes involved in oxidative stress, tissue homeostasis, neuronal maintenance, and retinal degeneration. Functional studies of one prioritized gene, *ARHGEF12*, revealed altered expression and phenotypes in disease-variant hiPSC-RGCs, supporting the utility of this framework for connecting POAG risk loci to disease mechanisms.

## Results

### Chromatin contact maps of hiPSC-RGCs and hTMCs

First, we sought to implicate candidate causal variant-effector gene pairs for the 46 POAG-associated loci previously identified by the POAAGG study.^38^ Chromatin contacts were captured using hiPSC-RGCs generated from individuals of African ancestry (**Table S1;** Penn003i-661-4, Penn039i-63-1, Penn059i-555-1) and commercially available hTMCs with no known/ reported retinal diseases. Additionally, we generated RNA-seq data for both models to contextualize gene expression in these cells (**Fig. 1a**). These experiments were sequenced to depth, passing standards set by consortia such as ENCODE, with three biological replicates for ATAC-seq, RNA-seq, and hTMC promoter-focused Capture C, and two biological replicates for hiPSC-RGC Hi-C (**Fig. S1a**). While Hi-C is a genome-wide approach, Promoter-focused Capture C enriches for contacts specifically at promoters, which can contribute to differences in contacts detected. Libraries passed other QC metrics, such as the expected fragment size distribution and proportion of long-range cis-reads in the chromatin interaction data (**Fig. S1b,c**).

**Figure 1:**
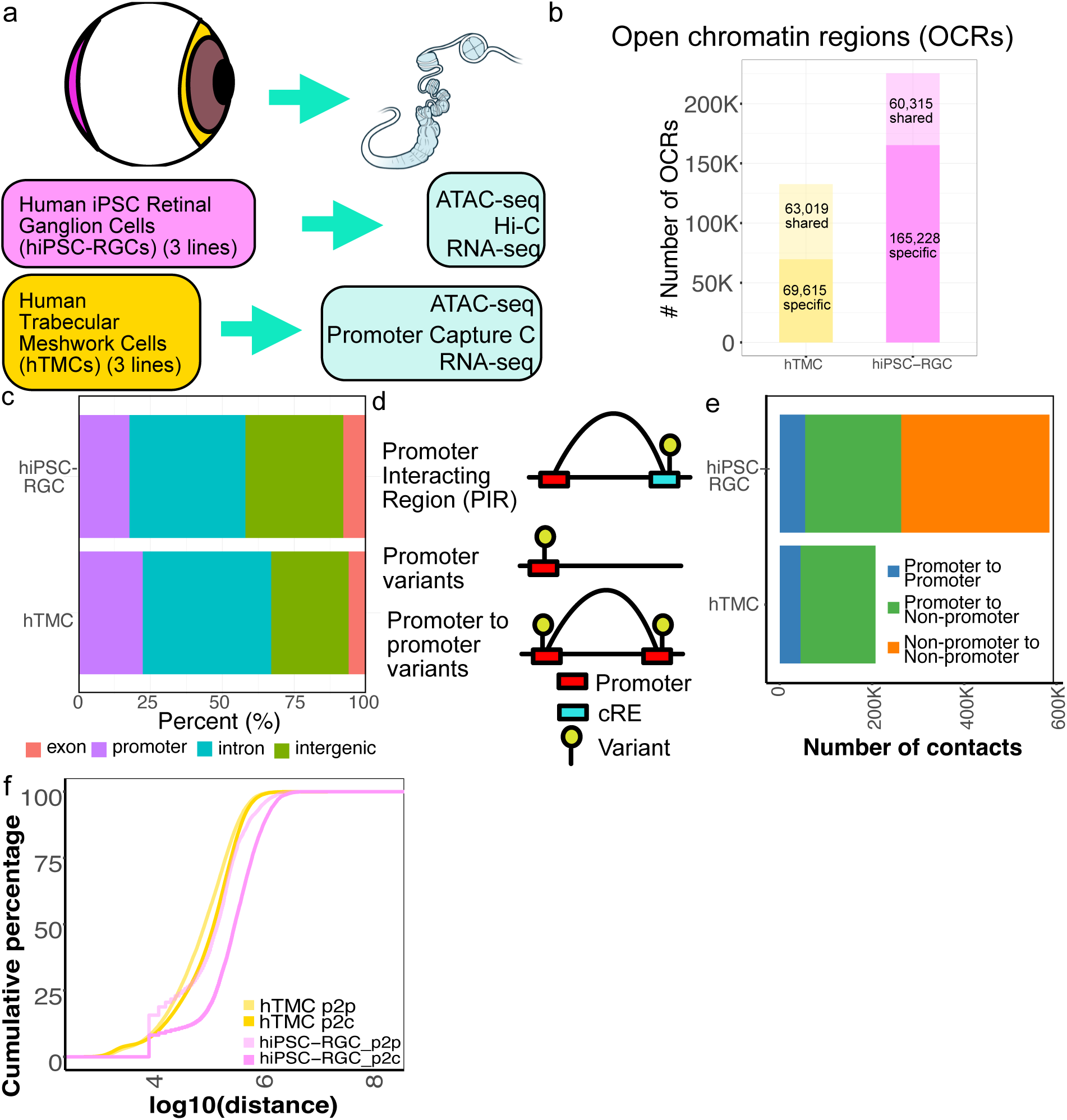
Constructing the cRE network of glaucoma-relevant cell types using ATAC-seq, RNA-seq, and chromatin conformation. (a) Schematic of the libraries constructed for the two cellular models of hiPSC-RGCs and hTMCs. While RNA-seq and ATAC-seq were performed on both hiPSC-RGCs and hTMC, chromatin conformation was characterized via Hi-C in hiPSC-RGCs and Promoter Focused Capture C in hTMCs. (b) The number of OCRs defined by ATAC-seq was specific to each cell type (yellow = hTMCs; pink = hiPSC-RGCs). OCRs were considered shared if there was a 1bp overlap with an OCR in the opposite cell type. (c) The proportion of OCRs falling into each category of genomic region annotations (exons, introns, intergenic, or promoter). (d) Schematic of cases for overlaying chromatin conformation with open chromatin data. Regions with significant interactions in either Promoter Focused Capture C or Hi-C are termed Promoter Interacting Regions (PIR). OCRs overlapping such regions are considered putative cREs for the targeted promoter. OCRs that overlay promoters (-1500bp/+500bp from a TSS) themselves are also considered as potential regulatory elements. Variants located in OCRs and nominated by these approaches are considered potentially regulatory. (e) The number of contacts per cell type that are between a promoter and a non-promoter region, between two promoters, or between two non-promoter regions (only found in Hi-C). (f) The cumulative distribution of the distance of significant chromatin interactions called in hTMCs and hiPSC-RGCs, stratified by either promoter to non-promoter (p2c) or promoter to promoter (p2p).

First, we focused on identifying open chromatin regions (OCRs) that may act as regulatory elements to influence gene expression. We identified 132,634 OCRs in hTMCs and 225,543 OCRs in hiPSC-RGCs that were reproducible in at least two replicates in each cell type. A subset of OCRs was shared in both cell types, with 47.5% of the OCRs found in hTMCs and 26.7% of the OCRs found in hiPSC-RGCs also detected in the reciprocal cell type (**Fig. 1b**). As expected, most OCRs were in non-coding regions such as introns and intergenic regions (**Fig. 1c**).

We then mapped these OCRs to effector genes using matched chromatin conformation data. We considered both proximal OCRs overlapping promoter regions themselves and OCRs overlapping distal promoter-interacting regions defined by Hi-C or Promoter-focused Capture C, as well as distal OCRs (**Fig. 1d**). The significant distal promoter interaction regions were considered using Chicago^63^ for promoter-focused hTMC Capture C data and the combination of Mustache^64^ and FitHiC2^65^ to call interactions in the hiPSC-RGC Hi-C data.^6,24^ We intersected the resulting loop calls with ATAC-seq peaks to define a set of promoter-interacting region OCRs (PIR-OCRs), which may function as long-distance cis-regulatory elements to influence gene expression. We further considered the set of OCRs located locally to a promoter (-1500/+500bp of the TSS) to be promoter OCRs.

A comparable number of promoter contacts were detected in both hiPSC-RGCs and hTMCs, with Hi-C also detecting many non-promoter contacts (**Fig. 1e**). A subset of these interactions involved one or more promoters (promoter-to-promoter interactions; p2p). Comparing the distance between these interactions, we detected that non-promoter to promoter interactions called by hiPSC-RGC Hi-C were longer than those called by Capture C in hTMCs. However, the same difference was not observed for promoter-to-promoter interactions across cell types (**Fig. 1f**). Overall, our approach identified 34,709 OCRs in interactions with 17,977 distinct genes in hTMCs and 39,496 OCRs contacting 19,174 genes in hiPSC-RGCs (**Fig. S1d,e**).

### Promoter-open chromatin interactions correlate with gene expression

Next, we validated the association of promoter contacts across OCRs in the context of gene regulation. We first identified the number of OCRs called per gene, which was a mean of 5.0 in hTMCs and 6.2 in hiPSC-RGCs (**Fig. 2a**). Similarly, we examined the mean number of genes connected per OCR, which was 2.6 in hTMCs and 2.5 in hiPSC-RGCs (**Fig. 2b**). Given that multiple genes may be implicated at a GWAS locus, we next sought to characterize the promoter-promoter and promoter-regulatory elements. We found 9.2% of genes in hiPSC-RGCs and 12.1% of genes in hTMCs were in promoter– promoter interactions with at least one other distinct gene. The number of connected OCRs per gene was positively correlated with gene expression across both cell types (**Fig. 2c**), suggesting that these interactions capture functional regulatory elements.

**Figure 2:**
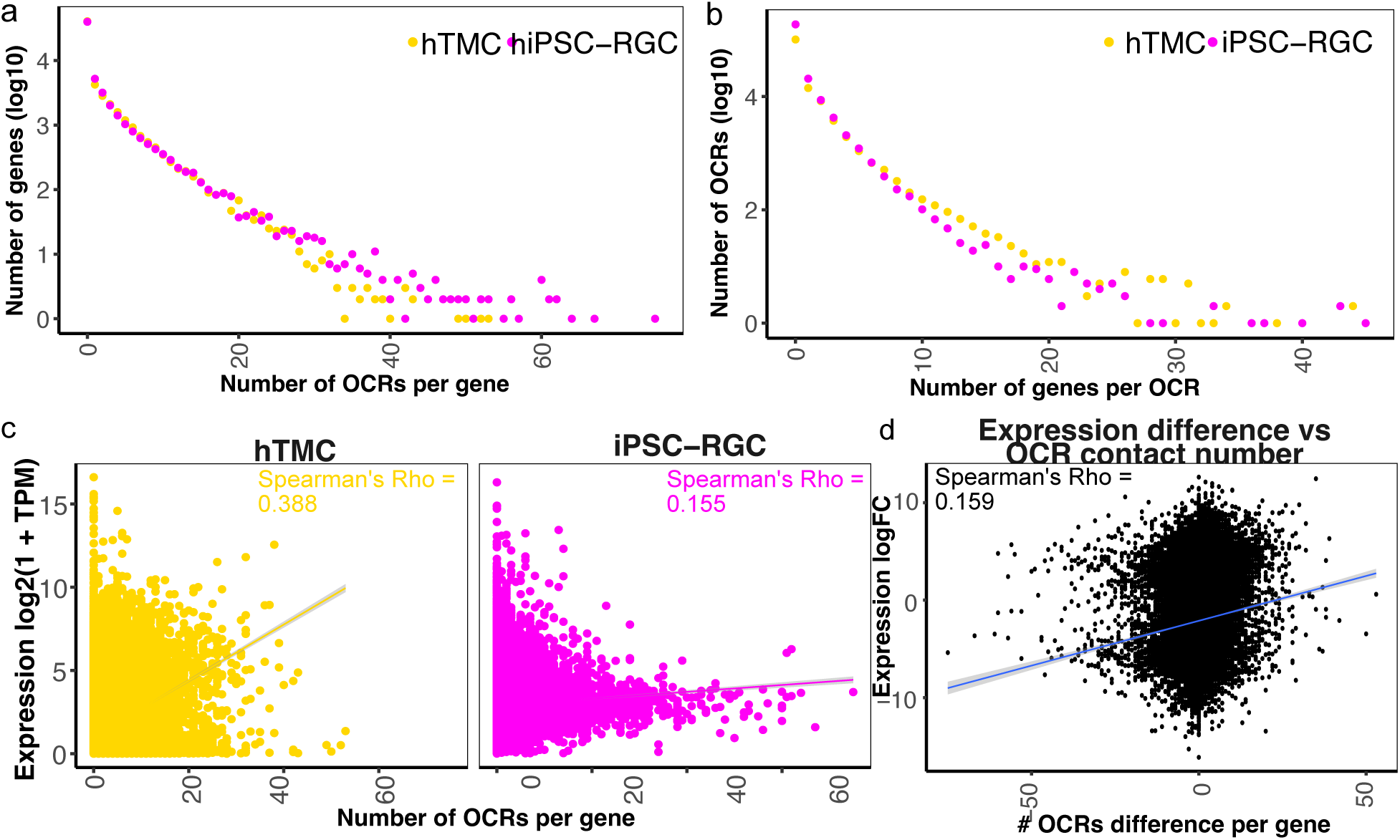
Promoter-connected OCRs are correlated with increased gene expression. (a) The number of OCRs per gene versus the log-scaled number of genes with the specified number of OCRs in each cell type (pink = hiPSC-RGC; yellow = hTMC). (b) The number of genes connected per log-scaled OCR for each cell type. (c) The number of OCRs contacting a gene promoter versus expression quantified in log2(1+TPM) for each cell type. d) The difference in expression against the difference in the number of OCRs predicted to be connected to each gene. In (c) and (d), the trend line from the linear association is shown with the shaded region representing 95% confidence interval.

Next, we examined whether the number of OCRs contacting promoters correlated with differences in gene expression between the two cell types. We identified differentially expressed genes between hiPSC-RGCs and hTMCs. Given that we expected large differences in expression between these two cellular models, we employed stringent criteria, as our primary goal in this analysis was to identify cellular marker genes. We observed 2,831 genes enriched in hTMCs and 12,316 genes enriched in hiPSC-RGCs (**Fig. S2a; Table S2**), which may reflect greater cellular diversity in generated hiPSC-RGCs when compared to directly harvested hTMCs. Reassuringly, known TMC and retinal neuron genes were detected, as the gene with the highest enrichment in hTMCs was *Fibronectin 1* (*FN1*), which is related to flow through the TMCs^66^, while in hiPSC-RGCs it was *Cerebellar Degeneration Related 1* (*CDR1*), which is involved in retinal neuron survival.^67^ Likewise, pathway enrichment analysis revealed enrichment of cytoskeletal, adhesion, and stress response pathways in hTMCs, which are all consistent with the clinical reduction of aqueous humor flow through hTMCs in POAG. Rhodopsin signaling, which plays an essential role in converting light into electrical pulses, displayed the strongest enrichment in hiPSC-RGCs (FDR = 0.08) (**Fig. S2b**). When we correlated the difference in the number of OCR contacts with gene expression, we observed a modest positive correlation (Spearman’s Rho = 0.159, *p* < 2.2e-19; **Fig. 2d**). Although statistically significant, this modest correlation may be limited because chromatin accessibility does not distinguish between active enhancers and other mechanisms that regulate gene expression. Taken together, these results are consistent with the expectation that increased chromatin contacts are positively associated with gene expression and may contribute to differences in expression across cell types that correlate with clinical phenotypes in POAG.

### Heritability enrichment of ocular traits in regulatory elements

Next, to determine which traits were enriched in our datasets, we conducted partitioned linkage disequilibrium (LD) score regression to assess enrichment of polygenic signals within genomic annotations across several eye-related traits. We included recent studies on vision-relevant traits, such as myopia and age-related macular degeneration, using publicly available GWAS summary statistics in European populations. We detected significant enrichments for myopia^68^, POAG^41^, intraocular pressure (IOP)^69^, progressive vision loss^70^, and macular telangiectasia^71^ (**Fig. 3a; Table S3**). For most traits that were enriched in both cell types, the hTMCs displayed stronger enrichment compared to hiPSC-RGCs, except for macular telangiectasia, a retinal disease.^71^ We did not detect significant enrichment in retinoschisis.^68^ Taken together, these analyses suggest that our datasets can serve as a useful resource for investigating the non-coding contribution of eye-relevant traits.

**Figure 3:**
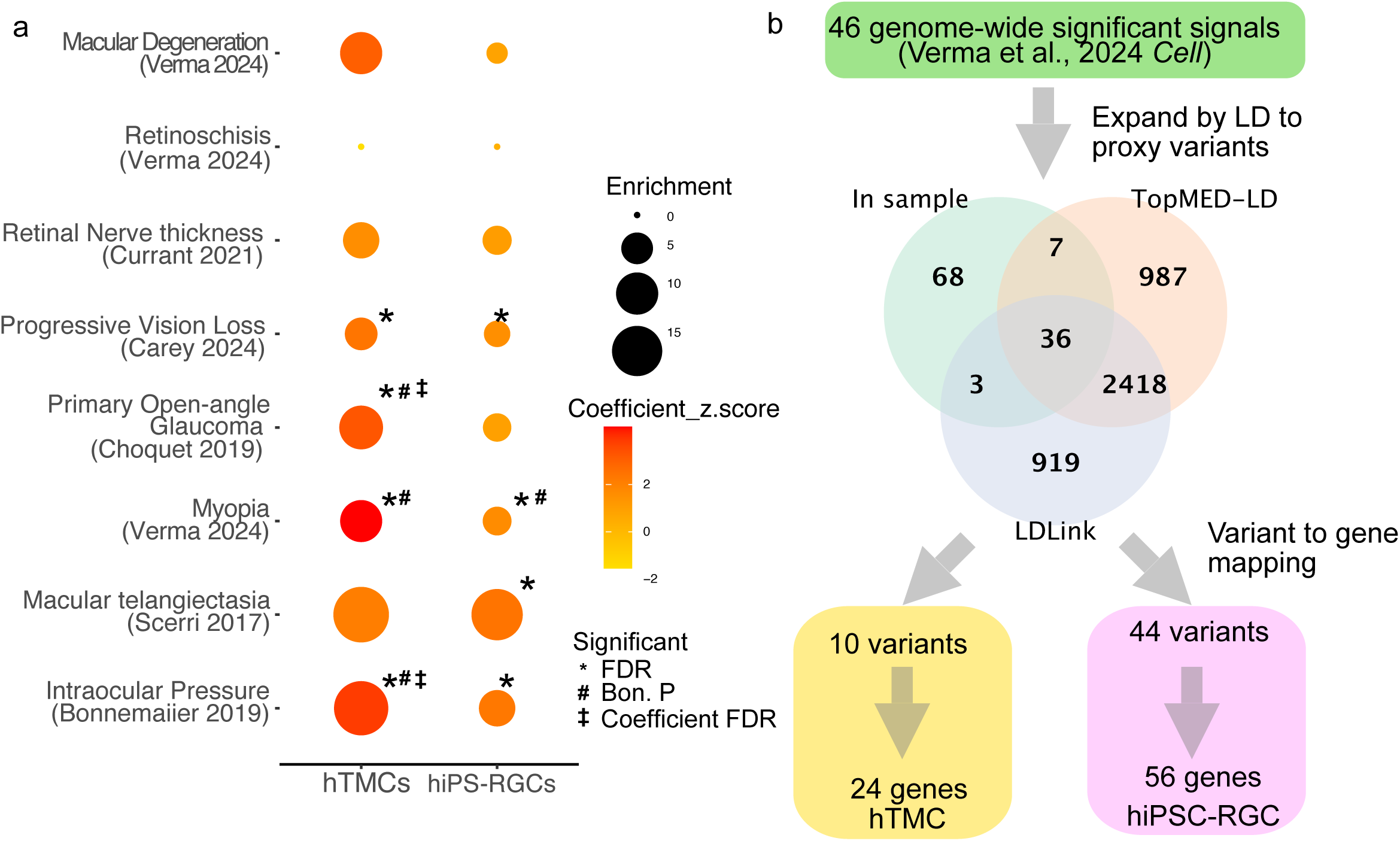
Dot plot depicting the results of partitioned LD score regression for hTMCs and hiPSC-RGCs across several European ancestry GWAS studies. Dot size depicts the size of enrichment and is colored by the coefficient z-score (higher corresponds to stronger association). *indicates the enrichment is significant following Bonferroni correction, # indicates the result is significant following FDR correction, and ‡ indicates significance of the baseline-adjusted cell type coefficient FDR. (b) To identify effector genes of risk loci identified in our African ancestry GWAS of POAG^38^, we expanded sentinel variants by LD (R2 > 0.6) using either the TOPMED-LD^74^ or LDLink^75^ reference databases. These variants were intersected with putative PIR-OCRs to identify the subset likely to be causal and their effector genes in hTMCs and hiPSC-RGCs.

While LD score regression is useful for adjusting for genetic confounding and heritability enrichment, the complexity of long-distance LD has limited its application for recently admixed groups, such as African Americans.^72^ Recently, LD score regression has been extended to accommodate these issues in covariate-adjusted LDSC^73^, which extends the LD window and uses principal components from the LD matrix as covariates for the regression. We performed partitioned LD score regression using our annotations with our recent African ancestry-based GWAS.^38^ While the GWAS sample size is slightly under the threshold for consistent signals used in the UK Biobank, we detected nominal significance with strong enrichment only in hTMCs (hTMC: Enrichment = 17.2, Unadjusted *p* = 0.044). This result, combined with the significant result in the European POAG study (**Fig. 3a**)^41^, motivated us to pursue variant-to-gene mapping for genome-wide significant POAG variants.^38^

### Variant-to-gene mapping of genome-wide significant POAG loci in African ancestry individuals

We next took 46 sentinel variants representing independent signals that achieved genome-wide significance in our prior GWAS of POAG in African ancestry individuals.^38^ We first expanded to proxy variants in strong linkage disequilibrium (LD; r^2^ > 0.6) using the sample matrix or TOPMED-LD^74^ and LDLink 1000G^75^ reference databases. This yielded 4,438 variants, with ∼56% agreement among two or more approaches (**Fig. 3b**). We then intersected these variants with OCRs showing significant chromatin contacts at a gene promoter, identifying 44 variants in hiPSC-RGCs that collectively interacted with 56 genes. Similarly, 10 variants in hTMCs were in contact with 24 genes (**Table S4**). Among the loci investigated, our approach nominated at least one gene for 20 loci (43% of signals).

From this list of putative effector genes, we prioritized variants based on: (1) overlap of findings from our prior orthogonal approaches, such as GWAS, and implication in other genetic studies of POAG (such as the *ARHGEF12* variant), (2) genes involved in neuronal function (*UCHL1, TUBB2B* and its associated lincRNA *LINC02525*), (3) genes involved in other retinal degeneration diseases (*CRB1* and *EXOSC2*), (4) antioxidant gene (*CAT*), (5) molecules involved in key cellular pathways of eye development such as Rho/Rho-associated kinase (ROCK) and Notch signaling pathways (*ARHGEF12*, *ROCK1*, *MIB2*, *MTA1* and its associated lincRNA *AL928654.1*).

We analyzed the mRNA expression of the selected genes in hTMCs isolated from human donor control eyes and POAG cases who underwent trabeculectomy (**Table S5**). First, we confirmed the authenticity of isolated hTMCs by assessing *myocilin* expression via qRT-PCR, which showed that dexamethasone-treated cells expressed significantly higher *MYOC* than untreated cells (**Fig. S3a-e**).^76^ When comparing mRNA expression of genes in hTMCs, we found that *ARHGEF12* (4 ± 0.13, *p* = 0.0001) and *ROCK1* (2.75 ± 0.12, *p* = 0.0001) were significantly elevated in hTMCs obtained from a POAG patient carrying a heterozygous SNP (rs11824032G>A). Similarly, *MIB2* was significantly elevated (1.72 ± 0.12, *p* = 0.0007) in POAG hTMCs with a homozygous SNP (rs4320726). However, there was no significant difference in the expression of *CAT* (**Fig. 4a-d**). Genotype information for the variants associated with the tested genes in each hTMC line is listed in **Table S6**.

**Figure 4:**
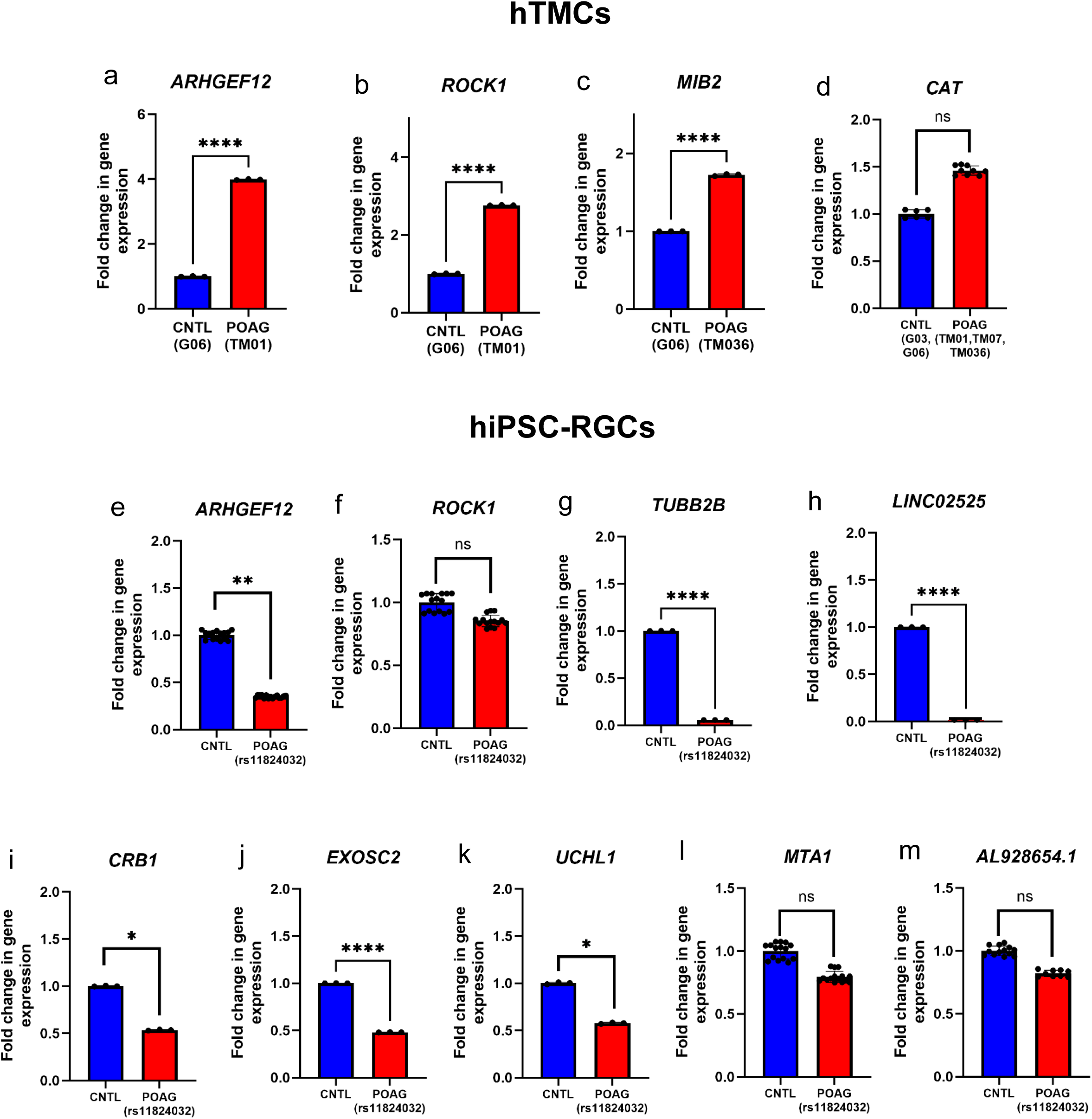
Quantitative mRNA expression analyses of prioritized genes implicated in hTMCs and hiPSC-RGCs. The quantitative expression profiles of the putativegenes using qRT-PCR, in cell lines with and without POAG. In hTMCs, genes include: (a) *ARHGEF12*, (b) *ROCK1*, (c) *MIB2*, and (d) *CAT*. In hiPSC-RGCs, genes include: (e) *ARHGEF12*, (f) *ROCK1*, (g) *TUBB2B*, (h) *LINC02525*, (i) *CRB1*, (j) *EXOSC2*, (k) *UCHL1*, (l) *MTA1*, and (m) *AL928654.1*. CNTL= cells from a visually unaffected control subject and POAG (rs11824032)= hiPSC-RGCs generated from a POAG subject with *ARHGEF12* SNP. Dots equal each technical replicate; *p<0.05, **p<0.01, ***p<0.001, ****p<0.0001.

Analysis of mRNA expression of genes in hiPSC-RGCs generated from a control line (9828) and a POAG line carrying the *ARHGEF12* variant (10030) (**Table S1**) showed that *ARHGEF12* was significantly downregulated (0.35 ± 0.05, *p* = 0.006) in the POAG hiPSC-RGCs. Similarly, *TUBB2B* (0.05 ± 0.07, *p* < 0.0001), *LINC02525* (0.015 ± 0.1, *p* < 0.0001), *EXOSC2* (0.48 ± 0.07, *p* < 0.0001), *UCHL1* (0.57 ± 0.15, *p* = 0.03), and *CRB1* (0.53 ± 0.15, *p* = 0.01) genes were all significantly downregulated in POAG hiPSC-RGCs compared to control cells. However, there was no significant difference in the expression of *ROCK1* and *MTA1*/ *AL928654.1* (**Fig. 4e-m**). Genotype information for each gene in the hiPSC-RGC lines is listed in **Table S7.**

### Functional validation of *ARHGEF12* variant in patient-derived hiPSC-RGCs

We initially identified *ARHGEF12* through fine mapping and colocalization analyses of POAG GWAS loci.^38^ We detected two variants (rs34002948 and rs11824032) corresponding to two independent signals (*r^2^* = 0) with contacts to promoters of this locus in hiPSC-RGCs (**Fig. 5a**). We predicted transcription factor motifs that could be either strengthened or disrupted by these variants (**Fig. 5b**). rs11824032 was predicted to disrupt several Kruppel-like zinc finger family motifs, while rs34002948 was predicted to strengthen affinity for several nuclear receptor family members. Among these, *KLF1* and *NR1I3* had the strongest expression in hiPSC-RGCs (**Fig. 5c**).

**Figure 5:**
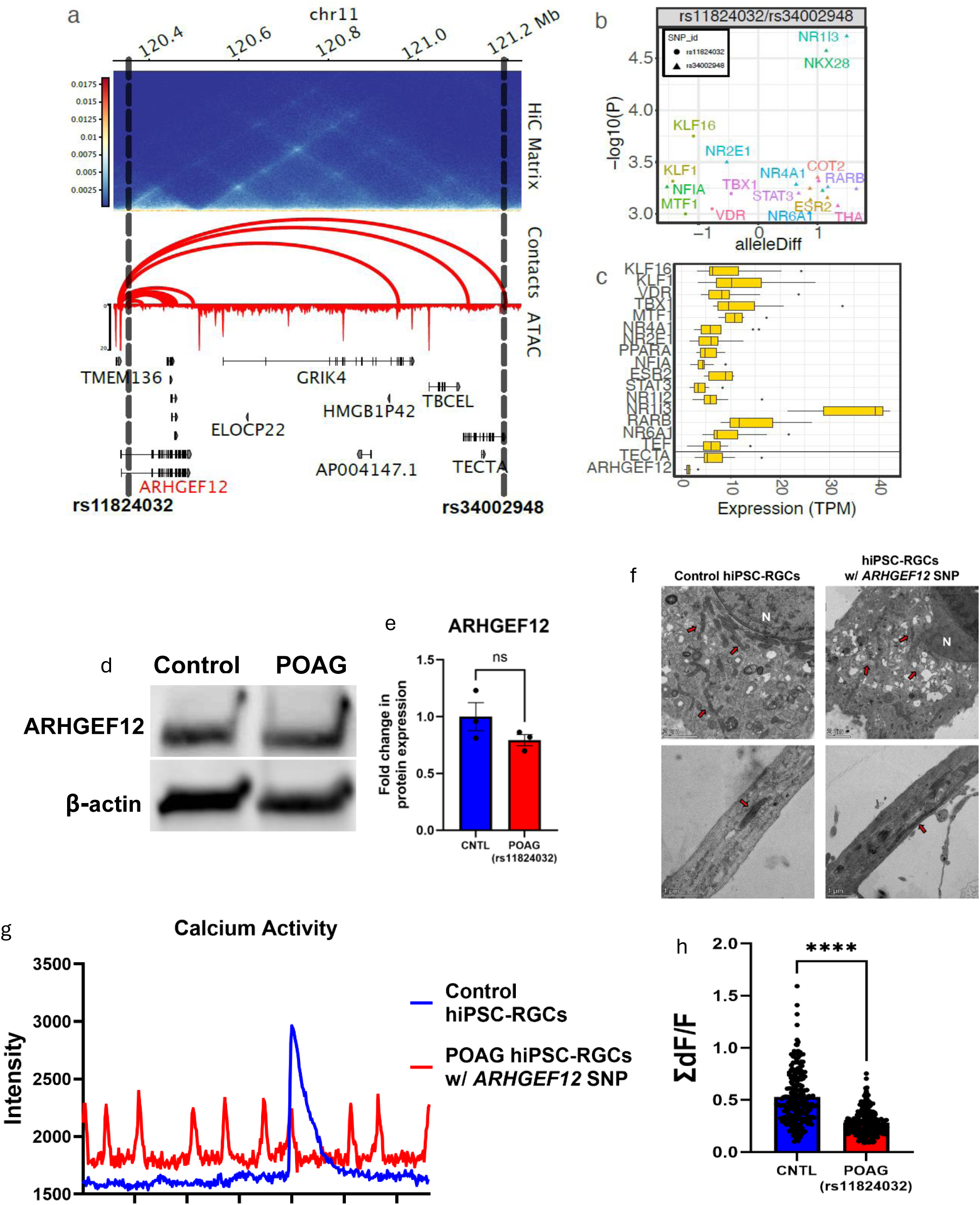
Nomination of *ARHGEF12* as a putative effector gene for African ancestry POAG in hiPSC-RGCs. (a) Genomic track showing contact frequency, promoter contacts, and chromatin accessibility for ARHGEF12. Two implicated SNPs, rs11824032 and rs34002948, are highlighted. (b) Motifs predicted to be disrupted by these variants (circles = rs11824032; triangles = rs34002948). The x-axis is the difference in information content (log scale) relative to the alternative/reference alleles, with the -log10 P value on the y-axis. Color corresponds to the transcription factor name. (c) The expression level of predicted transcription factors and implicated genes across each replicate of hiPSC-RGC in the *ARHGEF12* locus. (d) Western blots depicting ARHGEF12 protein and (e) quantification of the levels in control and POAG hiPSC-RGCs. (f) Transmission electron microscopy (TEM) images revealing ultrastructural analyses of morphological differences between control and *ARHGEF12* variant hiPSC-RGCs. Arrows pointing towards mitochondria and N: Nucleus. Scale bars as indicated. (g) Representative fluorescence responses from action potential firing in control and POAG hiPSC-RGCs. For simplicity, fluorescence from only one cell is shown for each condition. Note that the number of control RGCs exhibiting spontaneous Ca² responses is substantially greater than the number observed in POAG hiPSC-RGCs (see Videos 5a and 5b). (h) The total sum of all RGC amplitudes in control and *ARHGEF12* variant-expressing hiPSC-RGCs. CNTL= hiPSC-RGCs from a visually unaffected control subject and POAG (rs11824032) = hiPSC-RGCs generated from a POAG subject with the *ARHGEF12* SNP. Dots equal the number of cells quantified in each technical replicate; *p<0.05, **p<0.01, ***p<0.001, ****p<0.0001.

Together, multiple lines of evidence supported *ARHGEF12* as a candidate effector gene, including prior GWAS fine-mapping and colocalization results, chromatin conformation implicated *ARHGEF12* in both hTMCs and hiPSC-RGCs, and its established role in Rho/ROCK signaling. We note that expression in hiPSC-RGCs is lower than in hTMCs (**Fig. S2c**). Notably, the variant rs11824032 was in close proximity to the transcription start site (TSS) (<5 kb), whereas the other variant was implicated through a long-distance interaction (∼1 Mb) spanning several genes. Because this gene was moderately expressed (TPM =1.57) in control hiPSC-RGCs, we selected it for functional validation.

We confirmed a homozygous risk genotype for rs11824032G>A at the *ARHGEF12* gene in a POAAGG subject via Sanger sequencing (**Fig. S4b**) and selected an age-and sex-matched control (**Fig. S4a**). PBMCs were collected, reprogrammed into hiPSCs, and differentiated into RGCs using our 2D differentiation protocol.^77–79^ Day-35 FACS analysis showed *ARHGEF12*-variant hiPSC-RGCs expressed BRN3 (87%) and SNCG (92%) (**Fig. S4d**), while control hiPSC-RGCs also expressed BRN3 (96%) and SNCG (92%) (**Fig. S4c**).

Using western blot, we observed a trend toward decreased ARHGEF12 protein levels (0.79 ± 0.24) in hiPSC-RGCs derived from the POAG line (*n* = 3), but this difference was not statistically significant (**Fig. 5d-e**). However, ultrastructural analyses using transmission electron microscopy (TEM) revealed marked morphological differences between control and disease-variant hiPSC-RGCs. Control cells exhibited elongated mitochondria with an outer double membrane, intact cristae, dense matrix, and well-organized cytoplasm consistent with normal metabolic activity, whereas *ARHGEF12* variant cells displayed swollen, vacuolated mitochondria, cytoplasmic disorganization, and accumulation of autophagic vacuoles containing undigested material (**Fig. 5f**).^80^ These results highlight variant-linked cellular vulnerability.

To investigate the functional effects of the *ARHGEF12* variant on RGC activity, fluorescent Ca^2+^ indicators were used to indirectly measure neuronal responses of hiPSC-RGCs.^81–83^ Significant differences in fluorescence intensity were detected between the two hiPSC-RGC lines (*p* = 5.03e-14; **Fig. 5g**). Total RGC activity, which was expressed as the summed amplitudes of all Ca^2+^ responses calculated from the calcium activity intensity plots, was 53 ± 1.75% (*n* = 215) in control hiPSC-RGCs and 28 ± 0.82% (*n* = 213) in *ARHGEF12* variant-expressing hiPSC-RGCs (**Fig. 5h**). **Videos 5a** and **5b** illustrate representative Ca²⁺ signals in control and *ARHGEF12* variant hiPSC-RGCs, respectively (note that video frame rates were increased for presentation purposes).

## Discussion

POAG is highly heritable, yet known genes explain less than 10% of cases^84^ and over 30% of patients do not respond to current therapies.^85,86^ POAG pathogenesis remains unclear; it is reported that increased resistance to aqueous humor outflow by trabecular meshwork cells leads to elevated intraocular pressure, causing RGC death and optic nerve degeneration.^87^ While GWAS has identified many loci associated with POAG and its phenotypes, most are non-coding with small effect sizes. Here, we aimed to link POAG-associated SNPs to target genes and clarify the role of coding and non-coding variants in this disease. We integrated variants implicated by our large GWAS in the overaffected and understudied African ancestry population with gene expression analysis, accessible chromatin peaks, and 3D chromatin architecture maps in two glaucoma-specific cell types. Together, our analysis implicated 24 genes in hTMCs and 56 genes in hiPSC-RGCs with paired expression from the same donors. These cell type-specific variant lists will aid in understanding the gene-regulatory networks leading to vision loss in POAG patients.

We compared our hTMC Promoter-focused Capture C dataset from African ancestry-derived cells with a recently published Promoter Capture HiC dataset generated in hTMC samples from European ancestry individuals. Between the two datasets, fewer chromatin contacts (40,257) were reported in the European ancestry dataset compared to our dataset.^26^ Of these chromatin contacts, 60.0% overlapped with our interactions on both ends. The remaining differences in the datasets are likely driven by technical differences such as processing methods, restriction enzyme choice (Dpnll vs Arima kit), and differences in analysis such as using the geometric mean or hard filtering to handle replicates.^88,89^ Importantly, the substantial overlap supports the reproducibility of promoter interactions across studies, although future systematic comparisons will be needed to determine the extent to which chromatin architecture varies across ancestral backgrounds and experimental conditions.

We constructed a network of promoters and candidate cis-regulatory elements by linking promoters to ATAC-seq defined OCRs, generating a resource for interpreting GWAS variants associated with POAG and other ocular traits. Analysis of these networks revealed extensive regulatory complexity, with many promoters contacting multiple OCRs and OCRs shared across promoters. Regulatory elements in hTMCs and hiPSC-RGCs were enriched for relevant ocular diseases (**Fig. 3a**), with hTMCs showing stronger enrichment for heritability for most traits, which may be due to their role in regulating IOP. To further contextualize our findings, we compared the 24 implicated hTMC genes with published single-nucleus RNA sequencing data generated from control hTM tissue, finding that 11 were expressed in these cells. We have included their gene expression in specific cell types of the TM tissue (**Table S8**).^90^ While these datasets provide a valuable framework for variant-to-gene mapping, they were generated in a single cell state. Future studies investigating chromatin conformation in these cells under disease-relevant conditions, such as oxidative stress and glucocorticoids, may reveal additional regulatory interactions relevant to POAG pathogenesis.

We examined the expression of prioritized genes in both ocular cell lines, providing insight into the cellular basis of POAG risk. In hTMCs, prioritized genes point toward mechanisms involved in oxidative stress, tissue homeostasis, and regulation of aqueous humor outflow. Because TMCs are highly sensitive to oxidative stress,^103–105^ we further investigated the *CAT* variant, which is an antioxidant. Although prior studies have shown that serum samples collected from POAG patients exhibit low levels of catalase (CAT),^106^ we did not detect a significant difference in *CAT* expression between control and POAG hTMCs. In contrast, the *MIB2* gene showed significantly altered expression between control and POAG hTMCs with a homozygous SNP (rs4320726). This gene encodes the E3 ubiquitin ligase involved in Notch signaling, a pathway crucial for eye development. The Mouse Genome Informatics database reported that *Mib2-deficient* mice exhibit variable neural tube closure defects and abnormal eye morphology.^107,108^ This finding supports *MIB2* as a plausible effector gene at this locus and suggests that dysregulated Notch signaling may contribute to TM dysfunction in POAG.

Similarly, we investigated the mRNA expression of several genes of interest from our putative list in hiPSC-RGCs, showing that these genes converge on neuronal structure, synaptic function, and retinal degeneration pathways. The *TUBB2B* gene (*LINC02525*, its associated lincRNA) encodes ß-tubulin, a crucial building block of neuronal cell structure. Mutations in this gene have been reported to affect neuronal migration and axonal guidance, resulting in severe malformations.^109,110^ In our study, *TUBB2B* and *LINC02525* mRNA expression were significantly downregulated in POAG hiPSC-RGCs, suggesting disruption of cytoskeletal and axonal maintenance pathways that are essential for RGC survival. Another group reported that NMDA-mediated excitotoxicity downregulates *ARHGEF12* and *TUBB2B* expression in neurons.^111^ Similarly, *UCHL1*, encoding a ubiquitin enzyme, is specifically expressed in neurons and is involved in maintaining synaptic function.^112,113^ The downregulation of this gene in POAG hiPSC-RGCs is consistent with impaired neuronal function in POAG hiPSC-RGCs. The reduced expression of *CRB1*^114–121^ and *EXOSC2*^122^, which have known roles in retinal degenerations such as retinitis pigmentosa and Leber congenital amaurosis, suggests that POAG-associated genetic risk may converge on pathways shared with other neurodegenerative retinal disorders. *Arhgef12* is reported to be a modifier gene variant that contributes to *Crb1*-associated retinal dystrophy in mice.^123^ In the case of *MTA1*/ *AL928654.1*, we detected contacts between rs187398392 and the *Metastasis-associated 1* (*MTA1)* gene and associated lncRNA (*AL928654.1*/*MTA1-DT*), which are both established cancer genes. While *MTA1* has not been implicated in POAG, it has a known role in retinal development through interactions with *SIX3*, which regulates the number of retinal cells during development.^124^ In this study, we did not detect a significant difference in *MTA1* and *AL928654.1* expression between control and POAG hiPSC-RGCs.

One of the most compelling candidate effector genes implicated in both cell types was *ARHGEF12*. Notably, this gene has been associated with POAG and related phenotypes by several orthogonal approaches. Previous GWAS reports revealed an association between the *ARHGEF12* locus and endophenotypes, such as vertical cup-to-disc ratio (CDR) and IOP at the following lead variants: rs58073046^42,52^, rs199800298^41^, rs2305013^42^, and 11:120357425.^91^ Other studies have suggested *ARHGEF12* as a potential causal gene in the POAG GWAS locus based on expression quantitative trait loci (eQTLs).^46,58^ eQTLs acting on *ARHGEF12* in thyroid, brain, and fibroblasts in GTEx and a skeletal muscle sQTL were reported to colocalize with IOP GWAS, and sQTL in muscle, brain, and skin colocalized with POAG GWAS.^58^ Fibroblasts are relevant to the TM cell type, and brain tissue may capture the neuronal activity of RGCs or other neuronal cells in the retina. Furthermore, our group previously identified genetic colocalization between the lead variant (rs11824032) mapping to *ARHGEF12* and baseline CDR.^38^ Our variant-to-gene mapping results implicated rs11824032 as a putative causal SNP targeting its nearest gene, *ARHGEF12*. In addition, we also detected a long-range promoter contact to rs34002948, a variant associated with the “*TECTA*” locus (**Fig 5a; Table S4**). Given this gene’s nomination by two independent signals, both local and long-range to the gene body, we prioritized *ARHGEF12* for functional characterization in hiPSC-RGCs.^42,92^

EyeBrowse, an eye-centric genome browser, indicates that *ARHGEF12* is expressed in the TM, retina, and optic nerve.^93^ Furthermore, *Arhgef12* transcripts were detected in RGCs at P14 in mice.^94^ It is widely reported that this gene is a well-established component of the ABCA1-RhoA/ROCK signaling pathway, which regulates TM plasticity, aqueous humor outflow resistance, and subsequently increases IOP.^95–97^ But its functional role in the retina or RGCs remains poorly understood. In our study, we observed significantly elevated expression of *ARHGEF12* transcripts in POAG hTMCs compared to control hTMCs, but significantly downregulated expression in POAG hiPSC-RGCs, representing a novel finding. A similar trend was observed at the protein level, where we observed that POAG hiPSC-RGCs produced lower levels of ARHGEF12 protein than control hiPSC-RGCs. The eQTL colocalization analysis suggested that increased expression of *ARHGEF12* in the thyroid is associated with increased IOP levels, while decreased expression of *ARHGEF12* in the brain and cultured fibroblasts is associated with increased IOP, similar to the gene expression changes found in our study.^58^ Since ARHGEF12 is part of the Rho/ROCK signaling pathway, we also investigated expression of its downstream mediators. We detected a significant overexpression of *ROCK1* transcripts in POAG hTMCs carrying a heterozygous SNP (rs11824032G>A) at the *ARHGEF12* gene. However, there was no significant difference in *ROCK1* expression between control and POAG hiPSC-RGCs. Together, these results suggest that *ARHGEF12* may contribute to POAG through distinct cell type-specific mechanisms, influencing aqueous humor outflow regulation in the TM while affecting neuronal function in RGCs.

To further evaluate RGC dysfunction associated with the *ARHGEF12* risk variant, we evaluated RGC pathophysiology in control and POAG hiPSC-RGCs carrying the *ARHGEF12* variant using calcium imaging. Because Ca²⁺ influx through voltage-gated Ca²⁺ channels is tightly coupled to action potential firing, dynamic intracellular Ca²⁺ changes provide an indirect yet robust measure of neuronal function in hiPSC-RGCs. We observed reduced spontaneous activity in the *ARHGEF12* variant cells, with fluorescence changes (ΔF/F) of 28 ± 0.82% (*n* = 213) compared to 53 ± 1.75% (*n* = 215) in control cells. For context, naïve RGCs in wild-type mouse retina exhibit ΔF/F₀ values in the range of ∼150–200%,^98^ underscoring both the reduced responsiveness of hiPSC-RGCs relative to native tissue and the additional impairment associated with the *ARHGEF12* variant. We also performed single-cell patch-clamp recordings to assess electrophysiological properties directly; however, these experiments yielded inconclusive results due to low firing rates and sampling bias.^99–102^ Together, these findings suggest that the *ARHGEF12* risk genotype is associated with impaired neuronal activity in patient-derived RGCs, supporting a role for this locus in the neurodegenerative aspects of POAG pathogenesis.

In conclusion, our variant-to-gene mapping approach identified SNP-to-gene relationships and elucidated how variants contribute to POAG pathogenesis. By integrating LD-expanded variants with ATAC-seq and Hi-C data, we implicated 24 genes in hTMCs and 56 in hiPSC-RGCs. Our findings demonstrate how cell type-specific regulatory maps can bridge the gap between GWAS loci and biological mechanisms, providing a framework for prioritizing candidate effector genes for future functional studies. This extensive variant-to-gene dataset will serve as a key resource for the POAG research community going forward.

## Limitations

Our studies were conducted in hiPSC-RGCs derived from a single POAG case homozygous for the *ARHGEF12* variant, alongside one matched control. Therefore, until these findings are validated in hiPSC-RGCs from 2-3 additional cases and controls, we cannot conclude from these experiments alone that the observed differences are attributable solely to the *ARHGEF12* risk allele rs11824032. Nevertheless, we used these preliminary findings as a foundation for our subsequent functional experiments to characterize *ARHGEF12* and other variants of interest.

The hiPSC-RGC analysis of the *ARHGEF12* variant is preliminary, as we did not investigate how the variant affects RGC function. Ca²⁺ results should also be interpreted cautiously, since high-intensity, prolonged light stimulation may introduce artifacts. Additionally, we measured only intracellular calcium, not the electrical activity of RGCs.

This study focused on the *ARHGEF12* variant rs11824032, so hiPSCs were generated from a POAG subject with the risk allele, without controlling for other variants. To address this, we have combined hiPSC-RGCs with CRISPR/Cas9 to assess mutation-specific effects. Using prime editing, we aimed to create isogenic hiPSC lines with a single genetic background. Bulk editing efficiency for rs11824032G>A is ∼30% in HEK293T cells (**Fig. S5a-b**) and ∼15% and ∼12% in the two hiPSC lines, IMR-90 and WTC-11, respectively (**Fig. S5c-d**). Functional follow-up studies on the CRISPR-edited hiPSC lines are currently ongoing. Furthermore, POAG is a complex polygenic disease. To assess its heterogeneous pathogenicity, pooled CRISPR screens could be used to introduce gRNA libraries into hiPSCs, enabling gene overexpression or knockout.^125^ These cells can then be differentiated into RGCs, and sequencing of integrated gRNAs may reveal genes involved in RGC function and POAG pathogenesis.

## Materials and Methods

### Human iPSC culture

Undifferentiated hiPSCs were generated and characterized at the University of Pennsylvania iPSC Core Facility, providing a comprehensive analysis of hiPSC characteristics.^126–128^ The University of Pennsylvania and the Children’s Hospital of Philadelphia Human Subjects Research Institutional Review Boards approved all human sample collection protocols consistent with the Declaration of Helsinki. All methods were performed in accordance with the University of Pennsylvania’s relevant research guidelines and regulations. Written informed consent was obtained from all human cell donors. The hiPSCs were derived from the peripheral blood mononuclear cells (PBMCs) of the African donors and maintained in StemMACs iPSC-Brew XF (Miltenyi Biotec, catalog #: 130-107-086, North Rhine-Westphalia, Germany) containing 50X supplement (Miltenyi Biotec, catalog #: 130-107-087, North Rhine-Westphalia, Germany), and Pen Strep (Life Technologies, catalog #: 10378-016, Carlsbad, CA, USA). Table S1 lists the demographics of the subjects from whom the hiPSCs were generated.

### Retinal progenitor cell (RPC) generation and conditions

hiPSCs were cultured on Matrigel-coated plates (Corning, catalog #: 354230, Corning, NY, USA) in 37°C, 5% O_2_, and 5% CO_2_ conditions. Cells were maintained until they reached 75% confluency. hiPSCs were then split and seeded in one well of a 6-well tissue culture dish (Corning, catalog #: 229106, Corning, NY, USA) coated with 1:100 diluted Matrigel growth factor. hiPSCs were maintained at 37 °C, 5% O_2_, and 5% CO_2_ until reaching 100% confluence then transferred to 37 °C, 5% CO_2_ overnight prior to induction. On day 0, iPSC-Brew media was changed to RPC induction media: DMEM/F12 (50:50; Corning, catalog #: 10-092-cm, Corning, NY, USA), 1% Pen Strep (Life Technologies, catalog #: 10378-016, Carlsbad, CA, USA), 1% Glutamine MAX (Life Technologies, catalog #: 35050-061, Carlsbad, CA, USA), 1% NEAA (Life Technologies, catalog #: 11140-050, Carlsbad, CA, USA), 0.1 mM 2-ME (Life Technologies, catalog #: 21985-023, Carlsbad, CA, USA), 2% B27 supplement (w/o vitamin A; Life Technologies, catalog #: 12587-010, Carlsbad, CA, USA), 1% N2 supplement (Life Technologies, catalog #: 17502-048, Carlsbad, CA, USA), containing 2 μM XAV939 (X) (Wnt inhibitor; R&D Systems, catalog #: 3748, Minneapolis, MN, USA), 10 μM SB431542 (SB) (TGF-β inhibitor; R&D Systems, catalog #: 1614, Minneapolis, MN, USA), 100 nM LDN193189 (L) (BMP inhibitor; R&D Systems, catalog #: 6053, Minneapolis, MN, USA), 10 mM nicotinamide (Sigma-Aldrich, catalog #: N0636, St. Louis, MO, USA), and 10 ng/mL IGF1 (R&D Systems, catalog #: 291-G1, Minneapolis, MN, USA). Cultures were fed daily for 4 days. On day 4, the culture medium was exchanged for RPC induction media containing 2 μM XAV939, 10 μM SB431542, 100 nM LDN193189, 10 ng/mL IGF-1, and 10 ng/mL bFGF. The media was changed daily for 21 days.^79^

### RGC differentiation

Before RGC differentiation, the medium was changed daily using iNS media containing 250 ng/mL Shh (R&D Systems, catalog #: 8908-SH, Minneapolis, MN, USA), 100 ng/mL FGF8 (R&D Systems, catalog #: 423-F8, Minneapolis, MN, USA) for 2 days. For RGC differentiation on day 24, cells were manually crossed into small clusters with Leibovitz’s medium containing 34 μM D-Glucose (Research Products International, catalog #: G32045-500, Mt. Prospect, IL, USA) using the crosshatching technique as previously described^129^, then re-plated at a density of 1.0 x 10_5_ cells/well of 6-well plate coated with 1:100 diluted Matrigel growth factor in RGCs induction media containing: 100 ng/mL Follistatin 300 (R&D Systems, catalog #: 669-FO, Minneapolis, MN, USA), 0.5 μM Cyclopamine (R&D Systems, catalog #: 1623/1, Minneapolis, MN, USA), 3 μM DAPT (Stemgent, catalog #: 04-0041, Cambridge, MA, USA), and 4.2 µM Y-27632 dihydrochloride (Rock inhibitor; R&D Systems, catalog #: 1254/1, Minneapolis, MN, USA). The media was changed 24 hours post-plating with RGC induction media containing 100 ng/mL Follistatin 300 and 3 μM DAPT daily for 2 days. From day 27, media was changed to RGC induction media containing: 3 μM DAPT, 10 µM Rock inhibitor (Y27632), 5 μM Forskolin (Selleckchem, catalog #: S2449, Houston, TX, USA), 400 μM N^6^,2′-O-Dibutyryladenosine 3′,5′-cyclic monophosphate sodium salt (cAMP; Sigma-Aldrich, catalog #: D0627, St. Louis, MO, USA), 40 ng/mL BDNF (R&D Systems, catalog #: 248-BDB, Minneapolis, MN, USA), 5 ng/mL NT4 (R&D systems, catalog #: 268-N4, Minneapolis, MN, USA), and 10 ng/mL CNTF (R&D systems, catalog #: 257-NT, Minneapolis, MN, USA). The media was changed every 2-3 days until day 36. After maturation (D36), the medium was exchanged every 3-4 days with RGC induction media containing 3 μM DAPT and 10 µM Rock inhibitor (Y27632).^79^

### hTMCs

For the structural genomic experiments, such as RNA-seq, ATAC-seq, and promoter-focused Capture C, primary trabecular meshwork cells were used, which were commercially obtained (ScienCell, catalog #: 6590, Carlsbad, CA, USA).

For the functional validation of the variants, hTM tissue strips were isolated either from whole globes or obtained from glaucoma patients during trabeculectomy surgery at the Scheie Eye Institute. hTM tissues isolated from post-mortem donor eyes were obtained from the National Disease Research Interchange (NDRI). Whole globes were bisected near the equator, and the iris, lens, and ciliary processes were gently removed from the anterior segment. The anterior segment was then divided into two half-wedges. Using fine-tipped forceps, hTM tissue was grasped at the cut edge of each wedge and gently pulled away. Each hTM strip was cut into three pieces. hTM tissue from whole globes and trabeculectomy (typically 0.5–1 mm) was incubated for 30 minutes in DMEM (Gibco, catalog #: 11885-084, Waltham, MA, USA) containing 10 mg/mL dispase (Gibco, catalog #: 17105-041, Waltham, MA, USA) and 10 mg/mL collagenase (Gibco, catalog #: 17104-019, Waltham, MA, USA) at 37°C. Tissues were washed once with DMEM, then placed onto gelatin-coated plates with coverslips. hTM pieces from whole globes were distributed into three wells of a 6-well plate, while surgical hTM tissue was placed in one well. Coverslips were positioned to secure the tissue, and DMEM supplemented with 10– 20% FBS (Cytiva Hyclone, catalog #: SH30910.03, Logan, UT, USA) and penicillin-streptomycin (Gibco, catalog #: 15140-122, Waltham, MA, USA) was applied. Culture media was replaced every five days. Once the hTMCs reached confluence. Cultures were expanded further as needed or cryopreserved for long-term storage.^76^

We validated the authenticity of the hTMCs by performing *MYOC* induction experiments with 100 nM dexamethasone (Sigma-Aldrich, catalog #: D4902, St. Louis, MO, USA), freshly added to the media every 2-3 days for two weeks, which significantly increased *MYOC* expression by qRT-PCR relative to untreated controls.^76^ For all our experiments, we used primary TM cultures from Passage 2 (P2) to Passage 4 (P4). Demographics of the patients from whom the hTMCs were isolated are listed in **Table S5**.

### Sanger sequencing

DNA was extracted from hTMCs, control, and *ARHGEF12* disease-variant hiPSC-RGCs using the Purelink genomic DNA mini kit (Invitrogen, catalog #: K1820-01, Waltham, MA, USA). DNA was quantified using the Qubit dsDNA BR assay kit (Invitrogen, catalog #: Q32853, Waltham, MA, USA) and Qubit 2.0 fluorometer. PCR was performed using the GoTag Green Master Mix (Promega, catalog #: M7122, Madison, WI, USA), and the primer sets are listed in **Table S9**. PCR products were purified using Gel and PCR cleanup kit (Macherey-Nagel, catalog #: 74069.250, Düren, Germany) and submitted for Sanger sequencing at the Penn Genomic and Sequencing Core. Sequencing data were analyzed using NCBI BLAST and the UGENE (Unipro) software.

### ATAC-seq in hTMCs and hiPSC-RGCs

A total of 50,000 to 100,000 cells were centrifuged at 500 xg for 5 min at 4°C. The cell pellet was washed with cold PBS and resuspended in 50 µL of cold lysis buffer (10 mM Tris-HCl, pH 7.4, 10 mM NaCl, 3 mM MgCl2, 0.1% NP-40/IGEPAL CA630) and immediately centrifuged at 550 xg for 10 min at 4°C. Nuclei were resuspended in the Nextera transposition reaction mix (25 μL 2x TD Buffer, 2.5 μL Nextera Tn5 transposase (Illumina, USA Cat #FC-121-1030), and 22.5 μL nuclease-free H2O) on ice, then incubated for 45 min at 37°C. The tagmented DNA was then purified using the Qiagen MinElute kit, eluted with 10.5 μL Elution Buffer (EB). Ten microliters of purified tagmented DNA were PCR amplified using Nextera primers for 12 cycles to generate each library. The PCR reaction was subsequently cleaned up using 1.5x AMPureXP beads (Agencourt, Beverly, MA, USA), and concentrations were measured by Qubit. Libraries were paired-end sequenced using an Illumina NovaSeq 6000 (CHOP 51 bp read length).

Image files were demultiplexed using bcf2fastq (v2.20.0, Illumina, San Diego, CA, USA). Briefly, pair-end reads from three biological replicates for each cell type were aligned to hg38 genome using bowtie2,^130^ and duplicate reads were removed from the alignment. Narrow peaks were called independently for each replicate using macs2 (-p 0.01 --nomodel --shift -75 --extsize 150 -B --SPMR --keep-dup all -callsummits),^131^ and ENCODE blacklist regions were removed from peaks in individual replicates. The reproducible peak set (ATAC-seq peaks called by MACS peaks called with q-value < 0.01) was used as the consensus OCRs for each cell type. ATAC-seq fragment distribution plots were generated using the fragSizeDist function from the Bioconductor package ATACseqQC (v 1.28.0)^132^ with the deduplicated bam files.

### Hi-C in hiPSC-RGC

For the hiPSC-RGCs, Hi-C libraries were prepared for two biological replicates using the Arima-HiC kit (Arima Genomics, catalog #: A410030, Carlsbad, CA, USA), according to the manufacturer’s protocols. Briefly, cells were crosslinked using formaldehyde. Crosslinked cells were then subject to the Arima-HiC protocol, which utilizes multiple restriction enzymes (targeting ^GATC and G^ANTC) to digest chromatin. Arima-HiC sequencing libraries were prepared by first shearing purified proximally ligated DNA and then size-selecting 200–600 bp DNA fragments using AmpureXP beads (Beckman Coulter, catalog #: A63882, Brea, California, USA). The size-selected fragments were then enriched using Enrichment Beads (provided in the Arima-HiC kit) and then converted into Illumina-compatible sequencing libraries with the Swift Accel-NGS 2S Plus DNA Library Kit (Swift, catalog #: 21024, Ann Arbor, MI, USA) and Swift 2S Indexing Kit (Swift, catalog #: 26148, Ann Arbor, MI, USA). The purified, PCR-amplified DNA underwent standard QC (qPCR, Bioanalyzer, and KAPA Library Quantification (Roche, catalog #: KK4824, Basel, Switzerland)) and was sequenced with unique single indexes on the Illumina NovaSeq 6000 Sequencing System at 2x51 base pairs. For these biological replicates, two technical replicates were performed to fill the appropriate S4 flow cell.

### Promoter Capture C in hTMCs

For the hTM library, 107 fixed cells were thawed at 37°C, followed by centrifugation at RT for 5 mins at 1845 xg. The cell pellet was resuspended in 1 mL of dH2O supplemented with 5 μL 200X protease inhibitor cocktail, incubated on ice for 10 minutes, and then centrifuged. Cell pellet was resuspended to a total volume of 650 μL in dH2O. 50 μL of cell suspension was set aside for pre-digestion QC, and the remaining sample was divided into 3 tubes. Both pre-digestion controls and samples underwent a pre-digestion incubation in a Thermomixer (BenchMark, Tempe, AZ, USA) with the addition of 0.3% SDS, 1x NEB DpnII restriction buffer, and dH_2_O for 1 hour at 37°C, shaking at 1,000 rpm. A 1.7% solution of Triton X-100 was added to each tube, and shaking was continued for another hour. After pre-digestion incubation, 10 μL of DpnII (NEB, 50 U/μL) was added to each sample tube only and continued shaking along with pre-digestion control until the end of the day. An additional 10 μL of DpnII was added to each digestion reaction, and the mixture was digested overnight. The next day, an additional 10 μL of DpnII was added, and the mixture continued to shake for another 2-3 hours. 100 uL of each digestion reaction was then removed, pooled into one 1.5 mL tube, and set aside for digestion efficiency QC. The remaining samples were heat-inactivated by incubating at 1000 rpm in a MultiTherm for 20 minutes at 65°C to inactivate the DpnII, followed by cooling on ice for an additional 20 minutes. Digested samples were ligated with 8 uL of T4 DNA ligase (HC ThermoFisher, 30 U/μL) and 1X ligase buffer at 1,000 rpm overnight at 16°C in a MultiTherm.

The next day, an additional 2 μL of T4 DNA ligase was spiked into each sample and incubated for another few hours. The ligated samples were then de-crosslinked overnight at 65°C with Proteinase K (20 mg/mL; Denville Scientific, Holliston, MA, USA) along with pre-digestion and digestion control. The following morning, both controls and ligated samples were incubated for 30 min at 37°C with RNase A (Millipore, Burlington, MA, USA), followed by phenol/chloroform extraction, ethanol precipitation at -20°C, the 3C libraries were centrifuged at 1000 xg for 45 min at 4°C to pellet the samples. The controls were centrifuged at 1845 xg. The pellets were resuspended in 70% ethanol and centrifuged as described above. The pellets of 3C libraries and controls were resuspended in 300 μL and 20 μL dH2O, respectively, and stored at -20°C. Sample concentrations were measured by Qubit. Digestion and ligation efficiencies were assessed by gel electrophoresis on a 0.9% agarose gel and by quantitative PCR (SYBR green; Thermo Fisher, Waltham, MA, USA).

Isolated DNA from 3C libraries was quantified using a Qubit fluorometer (Life Technologies, Carlsbad, CA, USA), and 10 μg of each library was sheared in dH_2_O using a QSonica Q800R to an average fragment size of 350bp. The QSonica settings used were 60% amplitude, 30 seconds on, 30 seconds off, with 2-minute intervals, for a total of 5 intervals at 4°C. After shearing, DNA was purified using AMPureXP beads (Agencourt, Beverly, MA, USA). DNA size was assessed using a Bioanalyzer 2100 with a DNA 1000 Chip (Agilent, Santa Clara, CA, USA), and DNA concentration was determined via Qubit. SureSelect XT library prep kits (Agilent, Santa Clara, CA, USA) were used to repair DNA ends and for adaptor ligation following the manufacturer’s protocol. Excess adaptors were removed using AMPureXP beads. Size and concentration were verified using a Bioanalyzer with a DNA 1000 Chip and a Qubit fluorometer before hybridization. One microgram of adaptor-ligated library was used as input for the SureSelect XT capture kit using the manufacturer’s protocol and our custom-designed 41K promoter Capture-C library. The quantity and quality of the captured library were assessed using a Bioanalyzer with a high-sensitivity DNA Chip and a Qubit fluorometer. SureSelect XT libraries were then paired-end sequenced on the Illumina NovaSeq 6000 platform (CHOP; 51 bp read length).

Hi-C and Promoter Capture C data were processed using the HiCUP pipeline (v0.7.4).^133^ Significant chromatin contacts were called with Chicago (Score > 5)^63^ for Promoter-focused Capture C data (Chicago Score > 5), and the combination of Mustache (FDR < 0.1)^64^ and FitHiC2 (FDR < 1x10-6)^65^ was used as performed previously.^6,24^

### RNA-seq in hTMCs and hiPSC-RGCs

RNA was isolated from ∼1 million of each cell type using Trizol Reagent (Invitrogen, Waltham, MA, USA), purified using the Directzol RNA Miniprep Kit (Zymo Research, Irvine, CA, USA), and depleted of contaminating genomic DNA using DNAse I. Purified RNA was checked for quality on a Bioanalyzer 2100 using the Nano RNA Chip, and samples with RIN > 7 were used for RNA-seq library preparation. RNA samples were depleted of rRNA using QIAseq Fastselect RNA removal kit (Qiagen, Hilden, Germany). Samples were then processed for library preparation using the SMARTer Stranded Total RNA Sample Prep Kit (Takara Bio, Kusatsu, Shiga, Japan) according to the manufacturer’s instructions. Briefly, the purified first-strand cDNA is amplified into RNA-seq libraries using SeqAmp DNA Polymerase and the Forward and Reverse PCR Primers from the Illumina Indexing Primer Set HT for Illumina. The quality and quantity of the libraries were assessed using the Agilent 2100 Bioanalyzer system and Qubit fluorometer (Life Technologies, Carlsbad, California, USA). Sequencing of the finalized libraries was performed on the NovaSeq 6000 platform at the CHOP Center for Spatial and Functional Genomics.

RNA-seq was processed using the STAR (v 2.7.9a) pipeline, aligned to hg38 with gencode V30 as the reference. Reads were counted using HTSeq (v2.0.2)^134^, and TPM values were calculated using the effective gene length.

Differential expression analysis was conducted using standard procedures in the edgeR framework.^135^ First gene counts were aggregated across technical replicates. Low-expressed genes were then filtered using the filterByExpr function with the default parameters. Normalization was performed with the trimmed-median of the means method implemented in calcNormFactors. Dispersions were estimated with replicate cell types as a group and parameter robust = TRUE. The model of 0 ∼ cell type was fitted using glmQLFit with parameter robust = TRUE and tested with glmQFLTest across the cell type contrast. P-values were adjusted using the Benjamini-Hochberg procedure. Given the extreme differences observed between cell types, we opted for a strict cutoff of abs(logFC) > 7 and FDR < 1e-6 to focus on the strongest differential marker genes. TPM values were calculated as previously described using the effect-gene length.^136^ Pathway enrichment was conducted using the hypergeometric test implemented in phyper in R (R Core Team, 2025).

### LD Score Regression

We checked our chromatin annotations for GWAS heritability enrichment using LD-score regression implemented in LDSC.^137^ Publicly available EUR summary statistics were processed using the munge_sumstats.py script provided with ldsc (https://github.com/bulik/ldsc). Annotation files were constructed using open chromatin regions defined for hTMCs or hiPSC-RGCs. For European GWAS studies, LD score regression (v 1.01)^137,138^ was run using the European 1000 Genomes Phase3 reference data and developer-provided baseline v2.2 reference files.^139^ Enrichment analyses were run with the default parameters, with the exception of –print-coefficients set. Tests were adjusted with Bonferroni correction and false discovery rate (FDR) using the R function p-adjust. The GWAS and summary information, such as percent of heritability in either cell type is reported in **Table S3**. Dot plots were generated using the ggplot2 package (v. 4.0.1). For AFR ancestry analyses, covariate-adjusted LD score regression^73^ was used to adjust enrichment estimates for recent admix ancestry by incorporating principal components calculated and using an expanded LD window. To construct the reference, the hg38 1000G plink reference files were retrieved from the plink website. Principal components were computed using the eigenstrat smartpca function^140^ with the default parameters except numoutevac: 10. These were used in conjunction with the bed file for the 52 binary baseline annotations, as well as hTMC and hiPSC-RGCs as above, to build annotations with --ld-wind-cm set to 20 as previously described.^73^ This was used as the annotation to calculate enrichment as above.

### Genomic Analyses

#### Comparative analyses for promoter contacts

TPM values were correlated with the number of promoter-connected OCRs using the Spearman correlation coefficient, implemented in the cor.test function with method=”speaman” from R (v4.4.0).

#### SNP selection and LD expansion

Sentinel SNPs were collected from our previous African ancestry GWAS.^38^ These were expanded to any SNPs in high LD (R^2^ > 0.6) with the signal from the LD matrix of the population, or the EUR/AFR variants from the 1000 genomes high coverage set from LDLink^75^ or TOPMED-LD.^74^

#### Variant to gene mapping

Using the expanded set of genome-wide significant GWAS sentinels and their proxy variants, we intersected variants with the set of OCRs that either overlapped known promoter regions (-1500/+500 bp of gencode V30 TSS) or OCR overlapping promoter interacting regions identified by Capture C/ Hi-C in either hiPSC-RGCs or hTMCs. Genomic coordinate overlaps were identified using the R package GenomicRanges (ver 1.56.2)^141^ against the hg38 genome reference.

#### Transcription factor motif disruption

We used motifbreakR (v.2.19.5)^142^ to predict transcription factors that may have disrupted binding by the highlighted variants (rs11824032, rs34002948, rs187398392). Position weight matrices from hocomoco v10^143^ were queried against the hg38 genome with the reference and alternative allele. Motifs with strong disruption were predicted using the motifBreakR function, with parameters: filterp = TRUE, threshold = 1e-3, method = “ic”, bkg = c(A=0.25, C=0.25, G=0.25, T=0.25), and BPPARAM=BiocParallel::bpparam(). P-values were calculated with the calculatePvalue() function. The TPM-adjusted expression values were then plotted to identify which of the putative regulators may act in hiPSC-RGCs.

### FACS analysis

FACS analysis was performed on hiPSC-RGC cultures to quantify the number of cells expressing RGC-specific markers. Cells were lifted using TrypLE Express (Invitrogen, catalog #: 12605-010, Waltham, MA, USA) and collected by centrifugation at 1600 rpm for 5 minutes at 4 °C. The pelleted cells were resuspended in 1X PBS supplemented with 0.5% bovine serum albumin and 0.1% sodium azide (FACS buffer). Cells were fixed in 4% paraformaldehyde (v/v) for 15 min at room temperature (RT), followed by permeabilization using 0.5% Tween-20 (v/v) for 10 min at RT. Cells were incubated with anti-BRN3 Alexa Fluor 594 (Santa Cruz, catalog #: sc-390780, Santa Cruz, CA, USA) and anti-SNCG Alexa Fluor 488 (Santa Cruz, catalog #: sc-65979, Santa Cruz, CA, USA). Stained cells were analyzed using LSRFortessa B in the Penn Cytomics and Cell Sorting Shared Resource Laboratory at the University of Pennsylvania (RRID:SCR_022376). The data were further analyzed using FACS software (FlowJo 10.8.1, Ashland, OR, USA).

### qRT-PCR

Total RNA was extracted from hTMCs and hiPSC-RGCs using RNeasy® Mini Kit (Qiagen, catalog #: 74104, Hilden, Germany). For each sample, 150 ng of total RNA was reverse transcribed using the SuperScript® III first-strand cDNA synthesis kit (Invitrogen, catalog #: 18080-051, Waltham, MA, USA). Amplified cDNA was quantified using Fast SYBR™ Green Master Mix (Applied Biosystems, catalog #: 4385612, Foster City, CA, USA) on a QuantStudio 6 Pro (Applied Biosystems, Foster City, CA, USA), and the primer sets are listed in **Table S10**. All results were normalized to *GAPDH* housekeeping control and were from three technical replicates for each experimental condition. The relative gene expression was compared to that of control cells to obtain normalized gene expression, which is represented as the mean ± SEM.

### Western blot

hiPSC-RGCs from control and *ARHGEF12*-variant lines were scraped off the dish and centrifuged at 1600 rpm for 5 minutes. The pellet was extracted using RIPA buffer (Cell Signaling, catalog #: 9806S, Danvers, MA, USA), which contained protease and phosphatase inhibitors (Roche, catalog #: 03115828001, Mannheim, Germany). Cells were sonicated (15 s X 3) and then incubated on ice for 20 minutes. After lysing the cells, cell debris was removed by centrifugation at 20,000 xg for 30 minutes, and the supernatant containing the cytoplasmic extract was transferred to a new tube. The protein content of the cell lysates was determined using the Qubit protein assay (Invitrogen, catalog #: Q33221, Waltham, MA, USA) with the Qubit 2.0 fluorometer and then normalized to ensure equal protein content. Protein extracts were analyzed using SDS-polyacrylamide gel electrophoresis with Mini-Protean precast gels (Bio-Rad, catalog #: 4561094, Hercules, CA, USA) and followed by a western blot. The samples were analyzed for ARHGEF12 protein (Life Technologies, catalog #: PA5143957, Carlsbad, CA, USA), and β-actin (Santa Cruz Biotechnology, catalog #: sc-47778, Santa Cruz, CA, USA) was used as the loading control. The experiments were repeated at least three times.

### Transmission electron microscopy (TEM)

hiPSC-RGCs from a healthy control line and a POAG-associated *ARHGEF12*-variant line were cultured on 14-mm glass coverslips placed in a standard 24-well plate. Cells were plated at a density of 1 × 10⁵ cells per well to achieve a uniform monolayer suitable for ultrastructural analysis. Cells were maintained under RGC differentiation conditions until they formed a healthy monolayer suitable for ultrastructural analysis.

Cells grown on coverslips were fixed with 0.1 M sodium cacodylate buffer (pH 7.4) containing 2.5% glutaraldehyde and 2.0% paraformaldehyde and incubated overnight at 4 °C. After thorough buffer washes, samples were post-fixed in 2.0% osmium tetroxide containing 1.5% potassium ferricyanide (K₃Fe(CN)₆) for 1 hour at room temperature, followed by rinsing with deionized water. En bloc staining was performed using 2% aqueous uranyl acetate.

The samples were then dehydrated through a graded ethanol series, infiltrated, and embedded in EMbed-812 resin (Electron Microscopy Sciences, Fort Washington, PA). Ultrathin sections were stained with uranyl acetate and Sato lead stain and examined using a Talos L120C transmission electron microscope equipped with a Ceta 16 M digital camera.

### Calcium imaging

Intracellular calcium within live hiPSC-RGCs was stained using BioTracker™ 609 Red Ca2+ AM Dye (Sigma-Aldrich, catalog #: SCT021, St. Louis, MO, USA). 8 µL of the dye (80 µM) was diluted in 902 µL of 1X PBS, along with 10 µL of Pluronic® F-127 (Sigma-Aldrich, catalog # 9003-11-6, St. Louis, MO, USA) and 80 µL of DMSO. hiPSC-RGCs were incubated with the diluted dye for 40 minutes at 37 °C. Images of cells were acquired using an Olympus Fluoview 1000 confocal laser-scanning microscope equipped with an Olympus XLPlan N at 25X magnification. Images were acquired at a resolution of 256 x 256 pixels, at ∼2 frames per second, and covered an area of 509 × 509 µm. During imaging, cells were continuously perfused with Ames’ medium equilibrated with a 5% CO₂ / 95% O_2_ gas mixture and maintained at 35 °C. Imaging parameters were optimized to capture large fields of view and monitor the activity of as many cells as possible, which necessarily resulted in slower scan rates (approximately 2 frames per second). Under these conditions, individual action potential-evoked Ca^2+^ transients cannot be temporally resolved; instead, firing activity is represented as a normalized fluorescence change (ΔF/F), where ΔF reflects the increase in fluorescence associated with action potential-evoked Ca^2+^ influx and, for short bursts of firing, with spike rate and spike number, population-average ΔF/F provides a useful metric for comparing functional activity across cell types. Fluorescence responses in somas and dendrites were calculated with custom, ScanImage-compatible algorithms in Matlab. Fluorescent structures were outlined by hand using the roipoly function (Signal Processing Toolbox; The MathWorks). The fluorescence intensity of a neuron is reported throughout as the average intensity of all pixels over its soma, including the nucleus. Data was analyzed with custom software (Matlab; The MathWorks). The mean time-dependent fluorescent signal from each RGC (*F*) was normalized by subtracting the mean pre-stimulus signal intensity *(F0)*, which is expressed as *ΔF = F – F0*. The normalized neuronal responses (*ΔF/F0*) for each RGC were then calculated.^98,144^

### Prime editing

#### Plasmid Constructs and Cloning

The pCMV-PEmax-P2A-GFP (Addgene plasmid # 180020) prime editor was used.^145^ pegRNA and nsgRNA’s were cloned into the pU6-tevopreq1-GG-acceptor (Addgene plasmid # 174038), using BsaI Golden Gate assembly (NEB).^146^ All oligos were ordered from Integrated DNA Technologies (IDT). pU6-Sp-pegRNA-ARHGEF12: pegRNA spacer, top strand oligonucleotide: 5’-CACCGTCTGATCTCCTCAGAGACTAGTTTC-3’; bottom strand oligonucleotide: 5’-CTCTGAAACTAGTCTCTGAGGAGATCAGAC-3’ and Installation 3’-extension, top strand oligonucleotide: 5’-GTCCTCCCACTGCATCCCTAGTCTCTGAGGAGAT-3’; bottom strand oligonucleotide: 5’-CGCGATCTCCTCAGAGACTAGGGATGCAGTGGGA-3’. pU6-ARHGEF12-nsgRNA-1: nsgRNA spacer 1, top strand oligonucleotide: 5’-CACCGCCTCCTCTGGGCTCCTCCTCGTTTC-3’; bottom strand oligonucleotide: 5’-CTCTGAAACGAGGAGGAGCCCAGAGGAGGC-3’. pU6-ARHGEF12-nsgRNA-2: nsgRNA spacer 2, top strand oligonucleotide: 5’-CACCGTCTTCTCACAGGCATCTCTGGTTTC-3’; bottom strand oligonucleotide: 5’-CTCTGAAACCAGAGATGCCTGTGAGAAGAC-3’. nsgRNA termination, top strand oligonucleotide: 5’-GTCCTTTTTTTTCGCGGTTCTATCTAG-3’; bottom strand oligonucleotide: 5’-CGCGCTAGATAGAACCGCGAAAAAAAA-3’, this is used in place of pegRNA 3’-extension when making nsgRNA using pU6-tevopreq1-GG-acceptor. Scaffold, top strand oligonucleotide: 5’-AGAGCTAAGAAATTAGCAAGTTGAAATAAGGCTAGTCCGTTATCAACTTGAAAAAGTGGGACCGAGTCG-3’; bottom strand oligonucleotide: 5’-GGACCGACTCGGTCCCACTTTTTCAAGTTGATAACGGACTAGCCTTATTTCAACTTGCTAATTTCTTAG-3’.

#### Cell culture and transfection of HEK293T cells

This was performed as previously described.^147^ HEK293T cells were cultured in DMEM (Thermo Fisher Scientific, Waltham, MA, USA) supplemented with 10% FBS. HEK239T cells were seeded, one day prior to transfection, at 50000 cells/well in a 24-well plate. The plasmid constitution for transfection is 1050ng:393.75ng:78.75ng of CMV-PEmax: U6-pegRNA: U6-ngRNA. Lipofectamine 2000 (Thermo Fisher, Waltham, MA, USA) and plasmid DNA were mixed at a 1:1 mass ratio. Cells were collected 72 hours post-transfection for DNA extraction and analysis. DNA extraction was performed as previously described.^148^ HEK293T cells were detached with trypsin, DPBS washed, and finally re-suspended in 30 µl of DPBS. HEK293T cells were subsequently incubated at 95°C for 20 mins, 4 ul of 20mg/ml Proteinase K (Promega, Fitchburg, WI, USA) was added, and a further incubation at 56°C for 1h followed by a final 30 min incubation at 95°C. This crude DNA extract was then used as template for PCR.

#### Cell culture and transfection of iPSCs IMR-90 or WTC-11

IMR-90 or WTC-11 cells were seeded at 200,000 cells/well in 12-well plates coated with Corning® Matrigel® Matrix (Corning, NY, USA) 24 hours prior to transfection. 1400 ng of PEmax-GFP, 525 ng of pU6-pegRNA, and 105 ng of nsgRNA2 were transfected to each well for a total of 2030 ng of DNA delivered to the cells via Lipofectamine Stem (Thermo Fisher Scientific, Waltham, MA, USA) transfection reagent. 72h post-transfection, cells were dissociated with TrypLE Select Enzyme (Thermo Fisher Scientific, Waltham, MA, USA) for five minutes at 37°C, resuspended in DPBS (Thermo Fisher Scientific, Waltham, MA, USA) and filtered through a 35 μm cell strainer (Thermo Fisher Scientific, Waltham, MA, USA) to obtain a single cell suspension. GFP-positive cells were sorted using fluorescence-activated cell sorting conducted by the Penn Cytomics and Cell Sorting Shared Resource Laboratory at the University of Pennsylvania (RRID: SCR_022376). GFP-positive cells were bulk sorted into 96-well plate at 1,000 cells per well for each triplicate. 7 days post-sorting, cells were dissociated with TrypLE Select Enzyme (Thermo Fisher Scientific, Waltham, MA, USA) and DNA was extracted using EasySep™ Total Nucleic Acid Extraction Kit (STEMCELL Technologies, Vancouver, BC, Canada). 50 ng of extracted DNA from both transfected cells and un-transfected cells was used as template for PCR amplification of the target locus.

#### Analysis of prime editing efficiency

The *ARHGEF12* rs11824032 SNV locus (Forward Primer: 5’-GTGGAGGTGGATGCCCAAC-3’, Reverse Primer: 5’-AATTGGGCCGACTGGGATTA-3’) was amplified using primers. The subsequent amplicon was processed by Genewiz for their Sanger sequencing service. Editing efficiency analysis was performed using EditCo’s ICE Analysis tool (ICE CRISPR Analysis. 2025. v3.0. EditCo Bio: https://ice.editco.bio/#/).

### Statistical analysis

For the quantitative gene and protein expression data, as well as *ΔF/F0*, all data are represented as mean ± SEM. Each experiment was conducted in triplicate for statistical analysis. Differences in gene transcript and protein expression, *ΔF/F0* between control and POAG hiPSC-RGCs and hTMCs were analyzed using the Student’s *t* test for two-group comparison by statistical software (GraphPad Prism 10.0; GraphPad Software, Inc., La Jolla, CA, USA). *p-values of < 0.05 were considered statistically significant.

## Authorship Contribution Statement

Conceptualization: V.V., M.C.P., S.F.A.G., and J.M.O.; methodology: V.V., M.C.P., S.F.A.G, and J.M.O.; software: M.C.P., Y.B., Y.Z., S.S.V., S.F.A.G., and J.M.O.; validation: V.V., M.C.P., S.F.A.G., and J.M.O.; formal analysis: V.V., M.C.P., S.F.A.G., and J.M.O.; investigation: V.V., M.C.P., J.A.P., S.N., J.H., M.H., M.L., A.A.E., H.V.G., V.R.M.C., B.L., A.M.B., A.G.R., Q.N.C., E.M.E., P.S.S., V.A.; resources: S.F.A.G. and J.M.O.; data curation: V.V., M.C.P., J.H., L.M., P.M.J.Q., S.F.A.G., and J.M.O.; writing – original draft: V.V., M.C.P., and R.J.S.; writing – review & editing: all authors; visualization: V.V., M.C.P., S.F.A.G., and J.M.O.; supervision: S.F.A.G. and J.M.O.; project administration: V.V., M.C.P., S.F.A.G., and J.M.O.; funding acquisition: S.FA.G. and J.M.O.

