## Supplemental Figures for "Variant-to-gene mapping in ocular cell types identifies candidate effector genes for primary open-angle glaucoma"

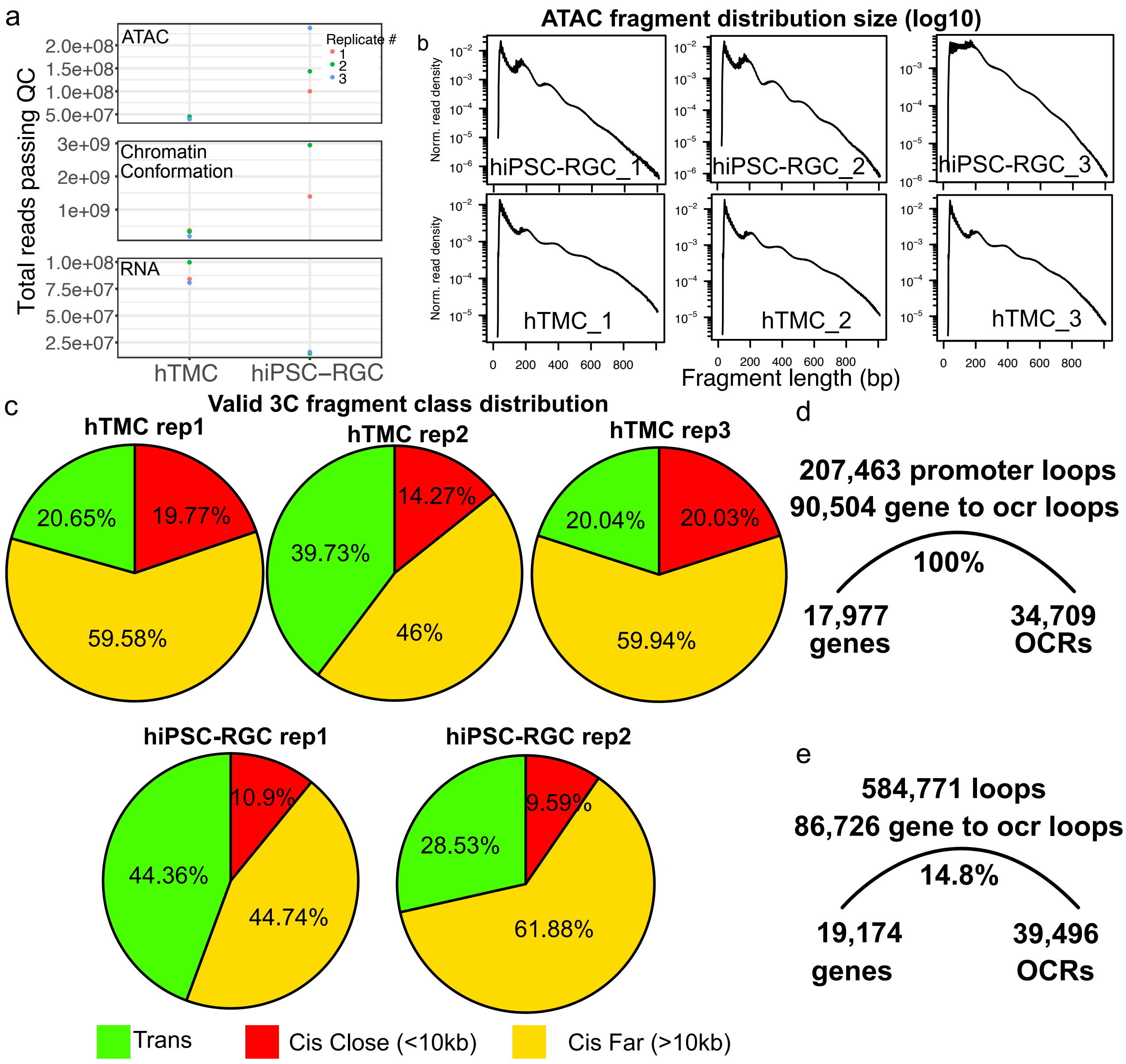


**Figure S1**: Quality control information for genome-wide assays. (a) Read counts for each biological replicate for ATAC-seq (both cell types), Promoter-focused Capture C (hTMCs), and Hi-C (hiPSC-RGC), and RNA-seq (both cell types). (b) Fragment length distribution for ATAC-seq, depicting mononucleosome and dinucleosome peaks. (c) Valid 3C fragment class distribution (cis-close, cis-far, trans). Summary of promoter loop connections between promoters and OCRs for (d) hTMC and (e) hiPSC-RGCs.

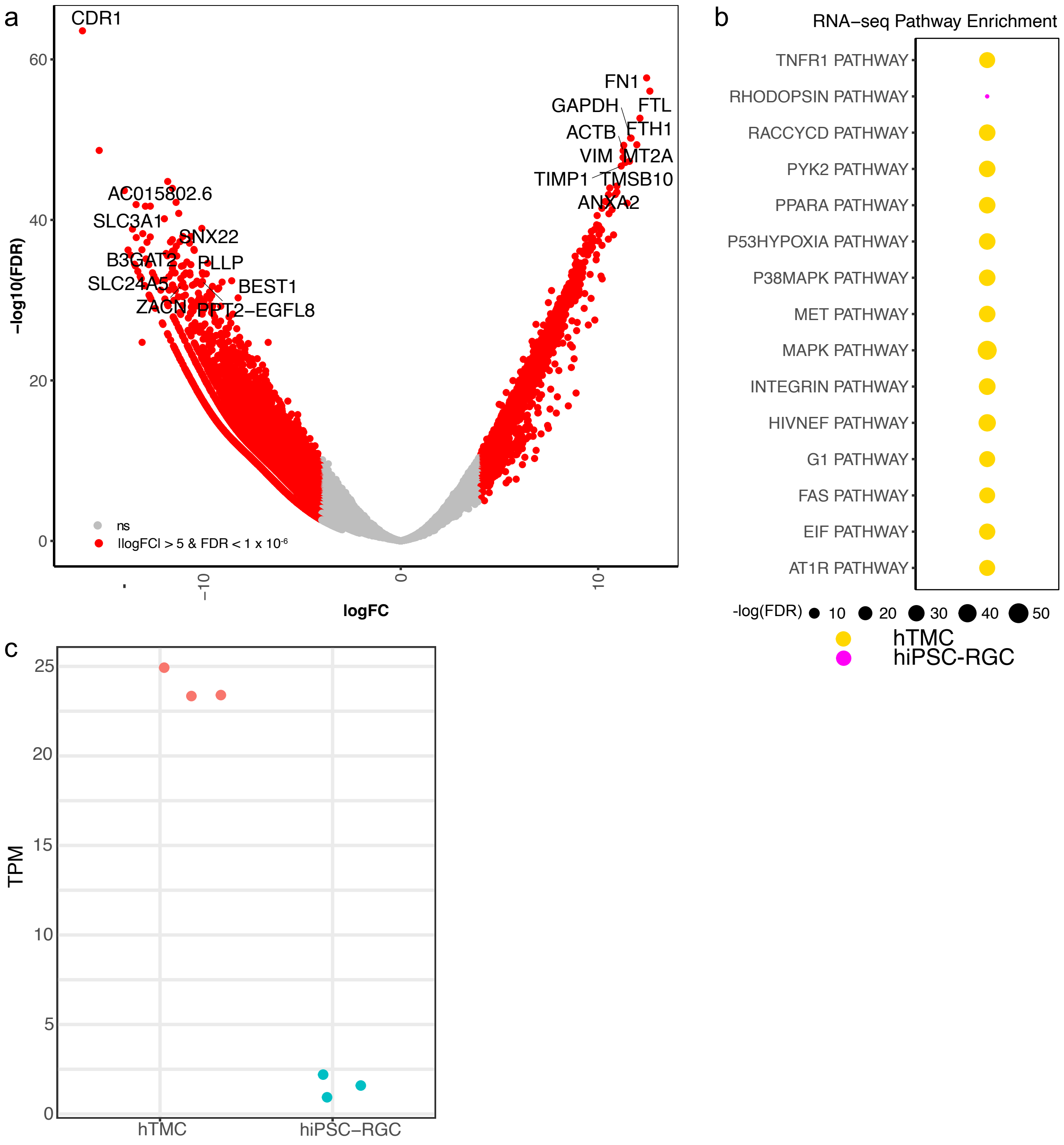


**Figure S2**: Differential gene expression analysis between hTMCs and hiPSC-RGCs. (a) Volcano plot with the x-axis depicting the log2 fold change between hTMCs (higher) and hiPSC-RGCs (lower). The y-axis depicts the -log10 transformed FDR value. (b) Dotplot depicts significantly enriched biological pathways from Camera analysis, the BioCarta collection on MSigDB140, with dot size corresponding to -log10(FDR-adjusted P value), while color indicates cell type (pink = hiPSC-RGC; yellow = hTMC). (c) TPM value for *ARHGEF12* in hiPSC-RGCs and hTMCs.

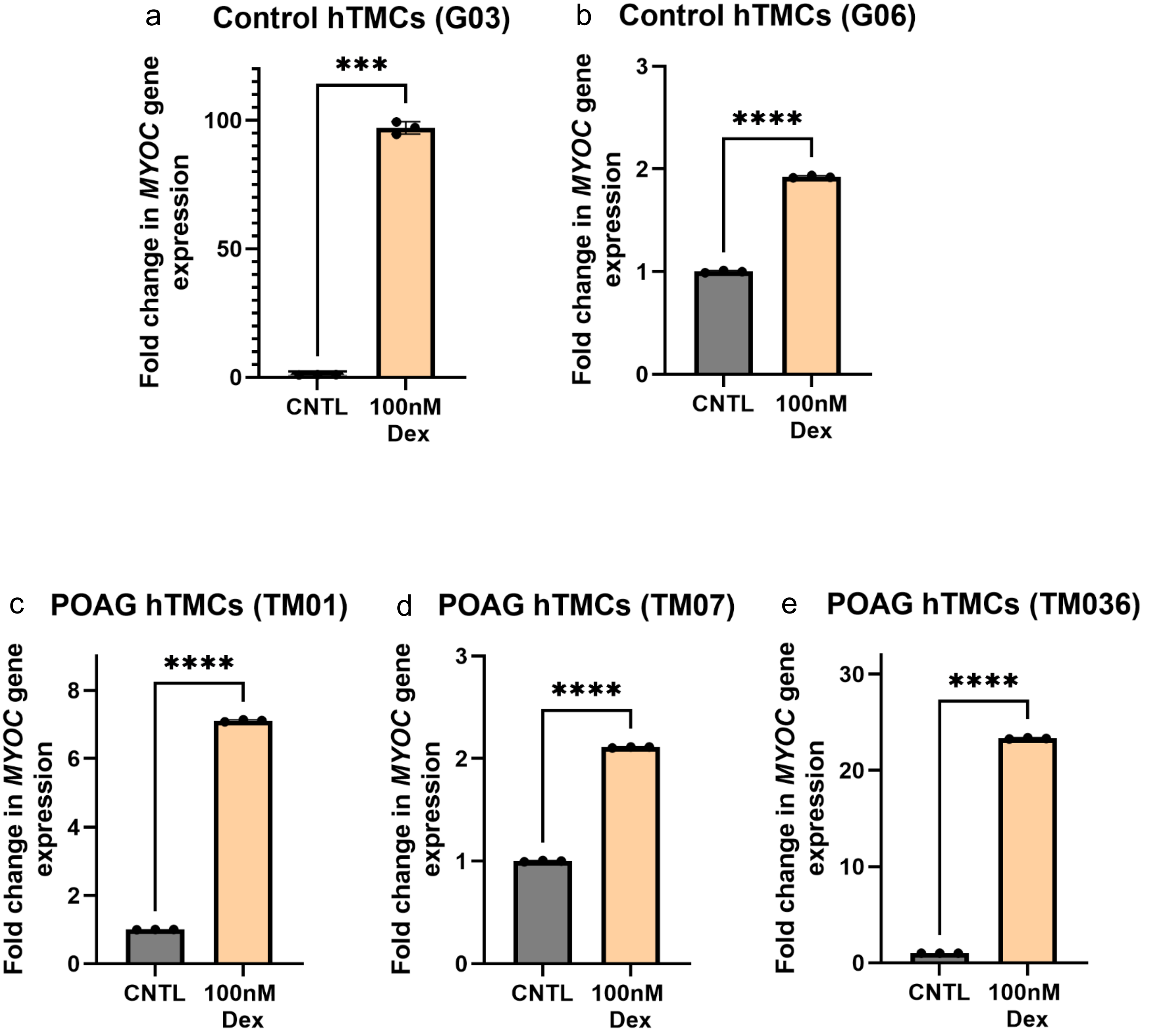


**Figure S3**: Characterization of the isolated hTM tissues by quantitative RT-PCR. We used qRT-PCR to confirm the presence of hTMCs isolated from human donor control eyes and POAG patients who underwent trabeculectomy. We determined that hTMCs treated with 100 nM of dexamethasone expressed significantly higher myocilin levels relative to untreated hTMCs in (a) G03, (b) G06, (c) TM01, (d) TM07, and (e) TM036. Dots equal technical replicates ($n$ = 3); *p<0.05, **p<0.01, ***p<0.001, ****p<0.0001.

a **Wild type hiPSC-RGCs**

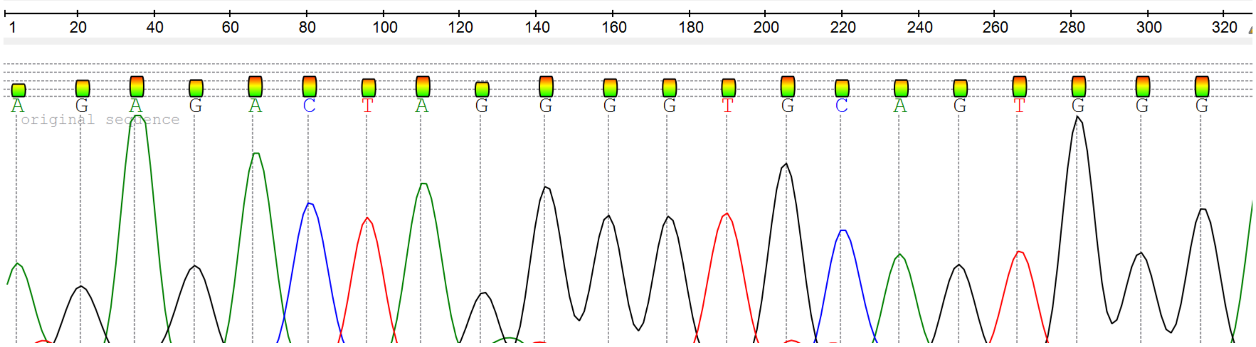


b ***ARHGEF12* hiPSC-RGCs (rs11824032G>A)**

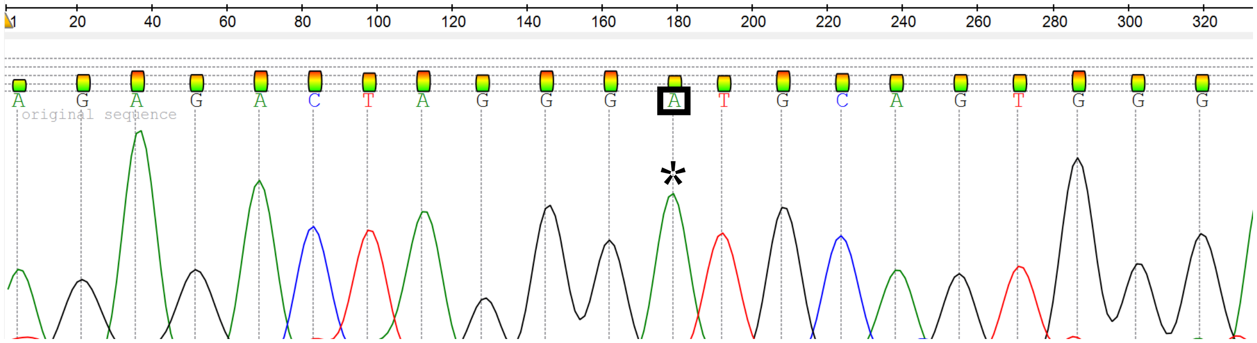


c **Control hiPSC-RGCs**

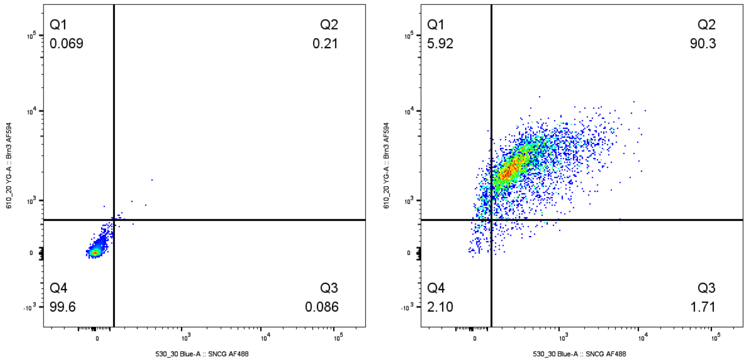


d ***ARHGEF12* hiPSC-RGCs**

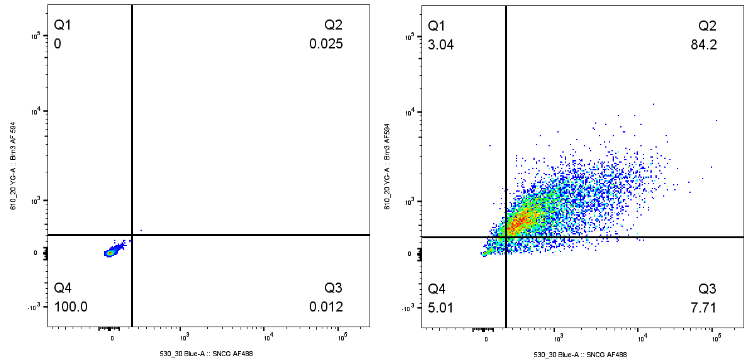


**Figure S4:** Characterization of the generated hiPSC-RGC lines. Sanger sequencing was performed to confirm the absence and presence of the rs11824032G>A variant in the *ARHGEF12* gene in (a) control hiPSCs and (b) POAG hiPSCs, respectively. FACS analysis was performed on the differentiated hiPSC-RGCs to confirm the presence of RGC-specific markers in (c) control hiPSC-RGCs and (d) POAG hiPSC-RGCs carrying the rs11824032G>A variant.

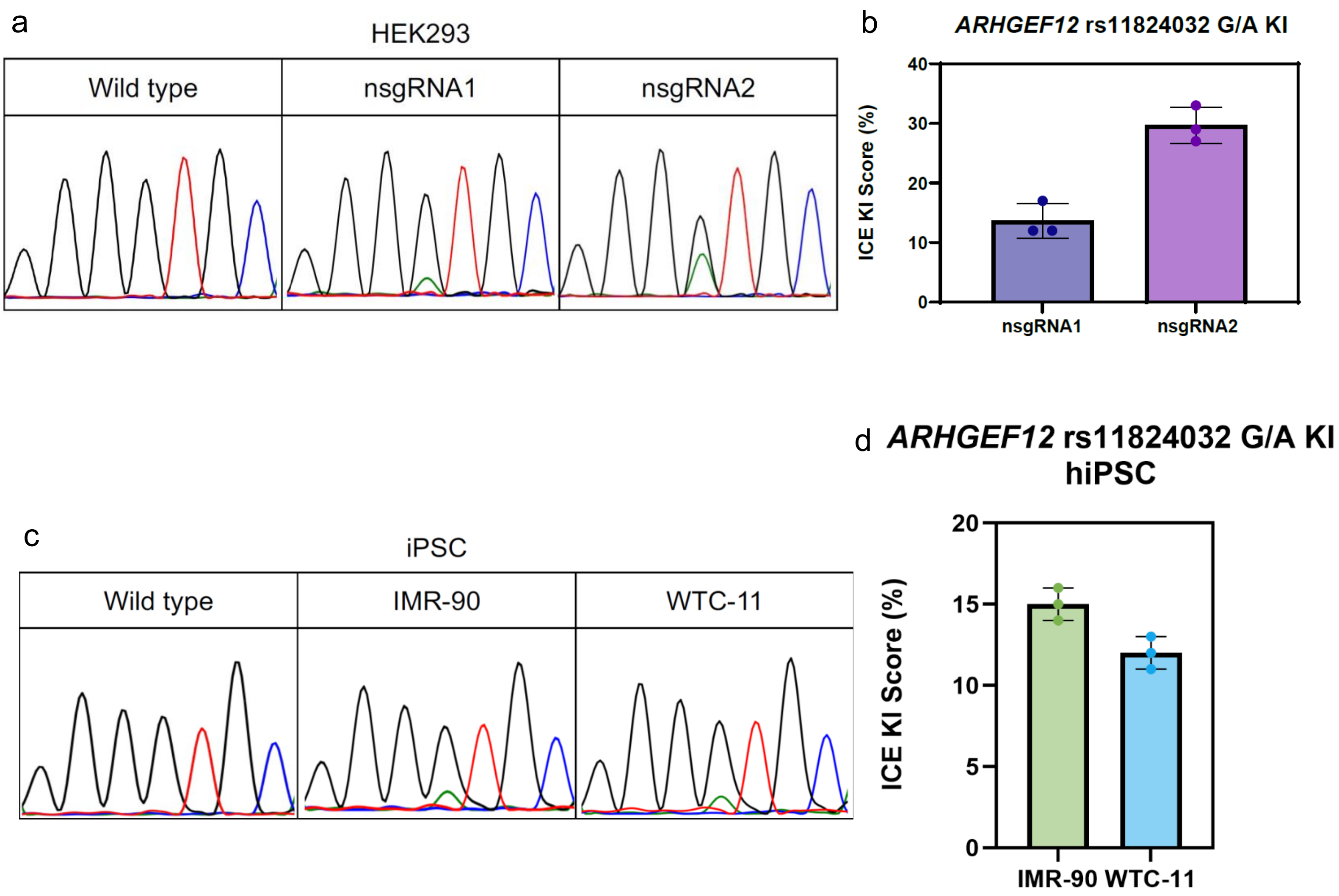


**Figure S5:** Prime editing of the *ARHGEF12* SNP in HEK293T cells and two hiPSC lines (IMR-90 and WTC-11). (a) Representative Sanger sequencing trace showing bulk editing of *ARHGEF12* rs11824032 G>A with nsgRNA1 or nsgRNA2. (b) Bulk prime editing in HEK293T cells for installation of the *ARHGEF12* rs11824032 G/A SNV. Nicking single-guide RNAs (nsgRNAs) 1 and 2 were used in conjunction with the pegRNA and the prime editor for prime editing. nsgRNA2 was used moving forward due to its higher editing efficiency as measured using Inference of CRISPR Edits (ICE) analysis. (c) Representative Sanger sequencing trace showing bulk editing of *ARHGEF12* rs11824032 G>A in IMR-90 and WTC-11. (d) Bulk editing efficiency of *ARHGEF12* rs11824032 G>A installation in two human iPSC lines, IMR-90 and WTC-11. Dots equal technical replicates ($n$ = 3).
